# Loss of the repressor REST affects progesterone receptor function and promotes uterine leiomyoma pathogenesis

**DOI:** 10.1101/2022.04.11.487902

**Authors:** Ashley S. Cloud, Faezeh Koohestani, Michelle M. McWilliams, Sornakala Ganeshkumar, Sumedha Gunewardena, Amanda Graham, Warren B. Nothnick, Vargheese M. Chennathukuzhi

## Abstract

Uterine Leiomyomas (UL) are benign tumors that arise in the myometrial layer of the uterus. The standard treatment option for UL is hysterectomy, although hormonal therapies such as selective progesterone receptor modulators are often used as temporary treatment options to reduce symptoms or slow the growth of tumors. However, since the pathogenesis of UL is poorly understood and most hormonal therapies are not based on UL-specific, divergent hormone signaling pathways, hallmarks that predict long-term efficacy and safety of pharmacotherapies remain largely undefined. In a previous study, we reported aberrant expression of REST/NRSF target genes activate UL growth due to the near ubiquitous loss of REST. Here, we show that ablation of the *Rest* gene in mouse uterus leads to UL phenotype and gene expression patterns analogous to UL, including altered estrogen and progesterone signaling pathways. We demonstrate that many of the genes dysregulated in UL harbor cis-regulatory elements bound by REST and progesterone receptor (PGR) adjacent to each other. Crucially, we identify an interaction between REST and PGR in healthy myometrium and present a putative mechanism for the dysregulation of progesterone responsive genes in UL ensuing the loss of REST. Using three *Rest* conditional knockout mouse lines, we provide a comprehensive picture of the impact loss of REST has in UL pathogenesis and in altering the response of UL to steroid hormones.

**Significance statement:** Ablation of *Rest* gene in the mouse uterus, modelling the loss REST in uterine fibroids, results in tumor formation and gene expression patterns analogous to human UL, including altered estrogen and progesterone receptor (PGR) pathways. The current study provides a putative mechanism for the aberrant function of PGR in UL.

## Introduction

Uterine fibroids (uterine leiomyoma, UL) are the most common reproductive tumors in women with an annual estimated cost of $5.9-34.3 billion in the US (1, 2). These tumors are found in the myometrial layer of the uterus and are characterized by increased proliferation of disordered smooth muscle cells (SMCs), enhanced estrogen sensitivity, and excessive deposition of extracellular matrix (ECM) (1, 3). Unfortunately, the heterogeneous nature of fibroids has limited our understanding of the pathogenesis of UL, and as a result hindered the development of effective pharmacotherapies (4).

Genetic, epigenetic, and chromosomal abnormalities as well as dysregulations in key signaling pathways involved in cell proliferation, apoptosis, ECM deposition and sex steroid hormone response have been known to play a role in UL development and growth (1, 4). Patterns of inherited genetic risk factors that predispose women to UL are extremely rare, whereas chromosomal abnormalities occur in 40-50% of UL tumors (5). In addition to chromosomal instability, a somatic mutation in mediator complex subunit 12 (*MED12*) has been reported to occur in up to 70% of UL, but not in the surrounding myometrium (6, 7). The molecular mechanisms that trigger these chromosomal events in the quiescent myometrial tissue are unknown.

Previously, we have shown that the aberrant widespread expression of GPR10 (PRLHR), led to uterine SMC proliferation through the activation of the PI3K/AKT-mTOR pathway (8). PRLHR is normally repressed in the periphery by the transcriptional repressor REST/NRSF (RE-1 silencing transcription factor/neuron-restrictive silencer factor) (9, 10). REST acts as a tumor suppressor in mammary epithelial cells and is shown to be downregulated in breast cancer, lung cancer, and colon cancer (11–14). We reported the loss of REST protein, but not mRNA, in UL promotes aberrant expression of REST target genes (8).

Ovarian steroid hormones, estrogen, and progesterone, are essential for UL tumor growth (15). An assortment of mechanisms that elevate estrogen signaling, including local estrogen production, increased expression of estrogen receptor 1 (ESR1) or PGR, and altered phosphorylation of ESR1 have been proposed as putative pathways that promote UL growth (16). Normally, progesterone suppresses ESR1 expression and cell proliferation in the myometrium through PGR (17). Molecular pathways that confer an aberrant mitogenic role for progesterone in UL are not well understood and may be key to developing long term pharmacotherapies for UL in the future (18).

We developed three UL-relevant conditional knockout mouse models to understand the role of *Rest* in the uterus. We demonstrate that the loss of REST in myometrium permits overexpression of its target genes leading to a UL phenotype consisting of SMC tumors and excess ECM production. Additionally, we provide evidence for the presence of conserved cis elements associated with REST and PGR adjacent to each other and a critical REST-PGR interaction in normal myometrium, which are essential for proper regulation of progesterone responsive genes. Absence of REST-PGR interaction due to the loss of REST alters steroid hormone response in UL. Based on the role that REST plays in steroid hormone sensitivity and UL tumorigenesis in the uterus, *Rest* cKO mouse models represent a crucial set of preclinical tools for the future development of pharmacotherapies for UL.

## Results

### Conditional Knockout of *Rest* in Mouse Uterus Leads to Leiomyoma Phenotype

Previously, we had shown that the loss of REST protein in leiomyoma patient samples leads to derepression of its target genes, including *GPR10* (8). To investigate the loss of REST *in vivo*, we developed a cKO of *Rest* in the mouse uterus since the conventional *Rest* KO is embryonically lethal (19). Chimeric founder mice were generated using embryonic stem cell clones in which exon 3 of *Rest* was floxed (Figure 1A, 1B) and homozygous reproductive tract specific *Rest^f/f^ Amhr2^+/Cre^* cKO mice were generated as described in Supplemental Methods. The *Rest^f/f^ Amhr2^+/Cre^* cKO mice had an increased uterine size compared to the control (Figure 1C & Supplemental Figure 1). Uteri of *Rest^f/f^ Amhr2^+/Cre^* mice were consistently hypertrophic, contained cystic glands, and had abnormal uterine morphology compared to control mice (Supplemental Figure 1). In addition, the cKO mice developed distinct fibroid tumors within the uterus that were not seen in control mice (Figure 1D, 1E). Immunostaining of mice uteri showed increase in myometrial thickness and excessive deposition of ECM components including collagen 3A1 (Supplemental Figure 2).

**Figure 1.**
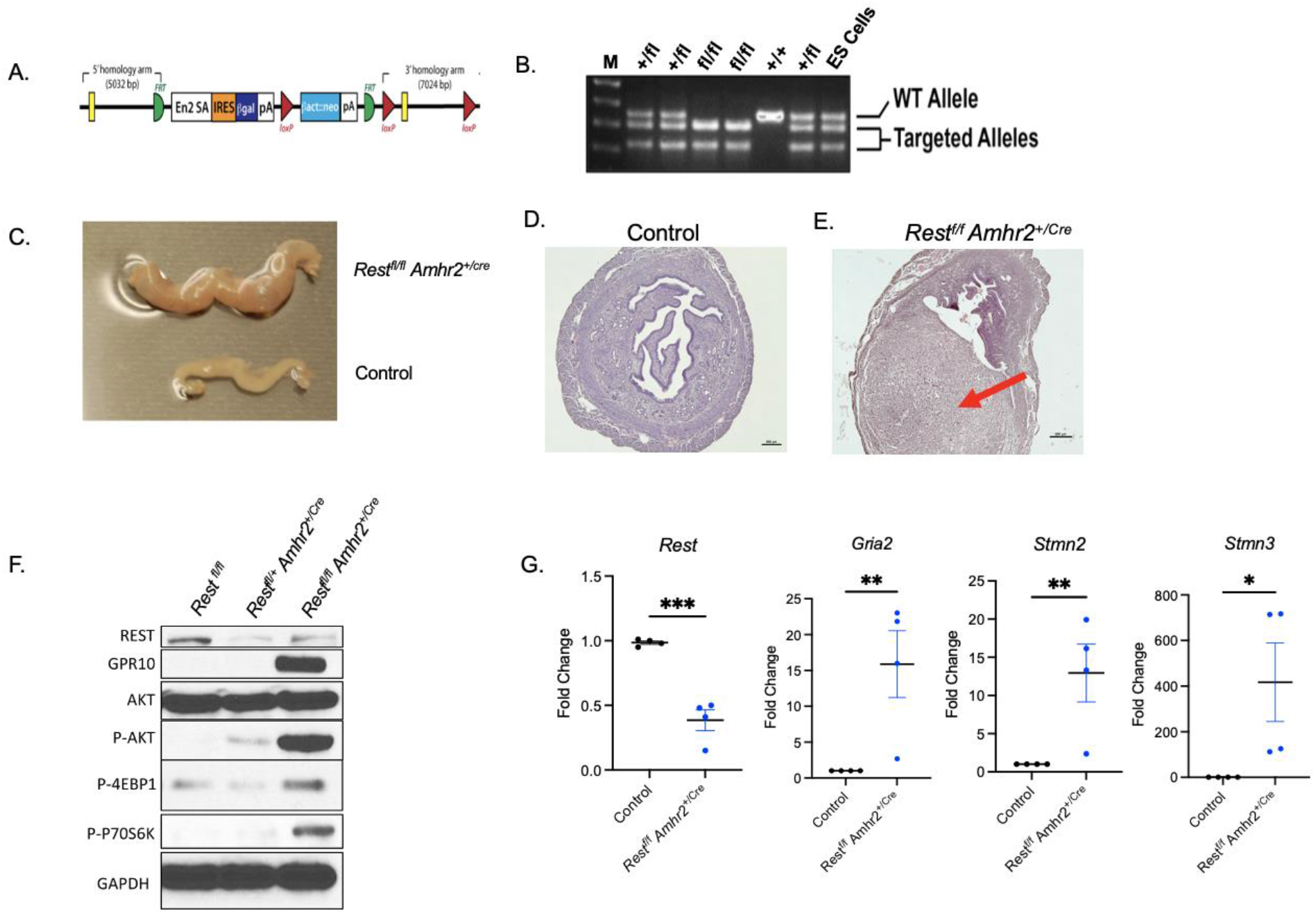
Loss of *Rest* under the *Amhr2* promoter displays a uterine leiomyoma phenotype. **(A)** *Rest^f/f^* targeting construct. **(B)** PCR genotyping of *Rest^fl/fl^* mice (lanes 4 & 5) *Rest* heterozygous mice (lanes 2, 3 & 7), and control mice (lane 6). (**C)** Representative image showing increased uterine size in 6-month-old *Rest^f/f^ Amhr2^+/Cre^* cKO mouse compared to control mouse during diestrus. **(D)** H&E stain of control mouse uterus. Scale bar: 500um. **(E)** H&E stain of *Rest^f/f^ Amhr2^+/Cre^* cKO with a tumor indicated by the red arrow. Scale bar: 500um. Images are at 4x magnification. **(F)** Western blot showing increased GPR10 expression and activation of downstream tumorigenic signaling proteins in total uterine tissue of *Rest^f/f^ Amhr2^+/Cre^* cKO mice compared to control and *Rest^fl/+^ Amhr2^+/Cre^* mice. GAPDH was used as a protein loading control. **(G)** Gene expression analysis of *Rest* and Rest targets, *Gria2*, *Stmn2*, and *Stmn3* in *Rest^f/f^ Amhr2^+/Cre^* compared to control (n=4). Error bars represent +SEM. Student’s T-test was performed, *P<0.05, **P<0.01, ***P=0.0001

Similar to the aberrant expression of GPR10 in UL specimen in which REST was lost (8), western blot analysis showed the ablation of *Rest* in the cKO mouse uterus resulted in an increase in GPR10 expression (Figure 1F). In addition, the cKO mouse uteri contained higher levels of phosphorylated forms of AKT, 4EBP1 and p70S6K, which are known mediators (8, 20) of the dysregulated PI3K/AKT-mTOR pathway in UL (Figure 1F). Quantitate RT-PCR confirmed significant decrease in *Rest* mRNA expression in the *Rest^f/f^ Amhr2^+/Cre^* cKO mice (Figure 1G). Additionally, REST target genes *Gria2, Stmn3* and *Stmn2* were all significantly overexpressed in the cKO mouse (Figure 1G), confirming our earlier findings that loss of REST leads to overexpression of its target genes, including *GRIA2, STMN2*, and *STMN3* (8). Many putative REST target genes, that have known roles in the uterus and significantly dysregulated in human UL, were also dysregulated in our *Rest^f/f^ Amhr2^+/Cre^* cKO mice, although species specific differences in gene regulation were present (Supplemental Table 1).

To further characterize the *Rest^f/f^ Amhr2^+/Cre^* cKO mice, gene expression profiles of the cKO and control mice in similar estrous cycle stage (diestrus) were analyzed using RNA sequencing (GSE178141). Ingenuity Pathway Analysis (IPA, Qiagen) revealed significant similarities in dysregulated gene expression profiles in human UL and *Rest^f/f^ Amhr2^+/Cre^* cKO mouse model, with most significant disease pathways in the knockout mouse uteri being analogous to leiomyomatosis, smooth muscle tumor, benign neoplasia, and leiomyoma (Table 1). Furthermore, the gene expression profile of the cKO mouse showed similarities to dermatological disorders (Table 1), hepatic fibrosis and retinoic acid signaling (Supplemental Figures 3A-C), diseases of reproductive and nervous system; all bearing overlapping cellular and molecular characteristics with UL (21–23). Lastly, the IPA confirmed REST to be downregulated and its network of targets to be significantly dysregulated as anticipated in the cKO model (Supplemental Figure 3D).

**Table 1.** Pathway analysis of functions or predicted diseases associated with dysregulated genes in UL and cKO mice. Disease predications made by Ingenuity Pathway Analysis software based off genes which were found to be dysregulated in both human UL (GEO dataset GSE13319; n=23) and RNA-sequencing results (GEO dataset GSE178141; n=3) of 4-month-old *Rest^f/f^ Amhr2^+/Cre^* cKO in diestrus.

| Diseases or functions | P-Value |
| --- | --- |
| Benign connective or soft tissue neoplasm | 4.85E-17 |
| Benign neoplasia | 1.74E-15 |
| Smooth muscle tumor | 5.24E-14 |
| Leiomyomatosis | 1.63E-13 |
| Benign neoplasm of female genital organ | 1.99E-13 |
| Connective or soft tissue tumor | 3.34E-12 |
| Uterine Leiomyoma | 8.04E-12 |
| Skin lesion | 3.65E-11 |

A highly relevant prediction of the IPA was how the loss of REST affected its upstream regulators, such as estrogen signaling (Supplemental Figure 3B). IPA identified β-estradiol as a highly significant upstream regulator of genes expressed in the *Rest^f/f^ Amhr2^+/Cre^* cKO, showing an activation of estrogen receptor signaling in the absence of REST. Indeed, the RNA sequencing on the cKO mouse uteri identified *Esr1* associated genes as significantly (*p*<0.05) dysregulated in these mice (Supplemental Figure 4). Moreover, we observed an increase in uterine size and hypertrophy in the cKO mice during diestrus when estrogen and progesterone were low and high, respectively (Figure 1C, Supplemental Table 3).

### Loss of *Rest* in Mouse Uterus Leads to Changes in Uterine Architecture

To study the impact of loss of REST in the uterus, we performed single cell RNA-sequencing using *Rest^f/f^PR^+/Cre^* cKO mice, instead of the *Rest^f/f^ Amhr2^+/Cre^* cKO mice, mainly due to the frequent embryonic lethality of *Rest cKO* due to leaky Cre expression under the *Amhr2* promoter as reported by others (24, 25). While the complete phenotypic characterization of the *Rest^f/f^PR^+/Cre^* is still ongoing, preliminary studies showed increase in the overall uterine size in 6-month-old mice compared to age-matched control uteri (Supplemental Figure 5). Additionally, the uteri of the *Rest^f/f^PR^+/Cre^* cKO mice were hypertrophic with cystic glands and abnormal lumen structure (Supplemental Figure 5), similar to *Rest^f/f^ Amhr2^+/Cre^* cKO mice.

Results from Seurat single cell RNA-sequencing data analysis (26) on uteri from 5-month-old *Rest^f/f^PR^+Cre^* cKO mice and littermate *Rest^f/f^* controls (GSE178141), which passed the quality control markers before being analyzed, identified 19 different clusters in the uterus based on their expression profiles (Figure 2A). Cell types were determined by gene expression profiles in each cluster using the SingleR software (27) and expert curation. A chi-square test of homogeneity comparing control and *Rest^f/f^PR^+/Cre^* cKO single cell RNA-sequencing cell counts revealed significant differences in population of neutrophils, epithelial, stromal, and myometrial cells (Figure 2B; Supplemental Figure 7). The results showing increased presence of neutrophils corroborates with reported upregulation of these cells in human UL (28). Based on conserved gene expression, cluster of cells representing smooth muscle cells (*Acta2, Cnn1, Myh11, Actg2, Myl9, Tpm2, Pcp4, Mylk, Tagln*, Supplemental Figure 8) as well as uterine stromal, myometrial fibroblast lineage (*Slco5a1, Col5a2, Col6a3, Dpt, Ddr2, Adamts4, Ifi205, Tgfb2, Cd34*, Supplemental Figure 9) were identified. Analysis of top TCA features (average cluster expression) of genes showed upregulation of ECM components in myometrial and stromal fibroblasts as well as smooth muscle cells (clusters 0 - 4, 8, and 17) in *Rest^f/f^ PR^+/Cre^* mice (Supplemental Figure 10).

**Figure 2.**
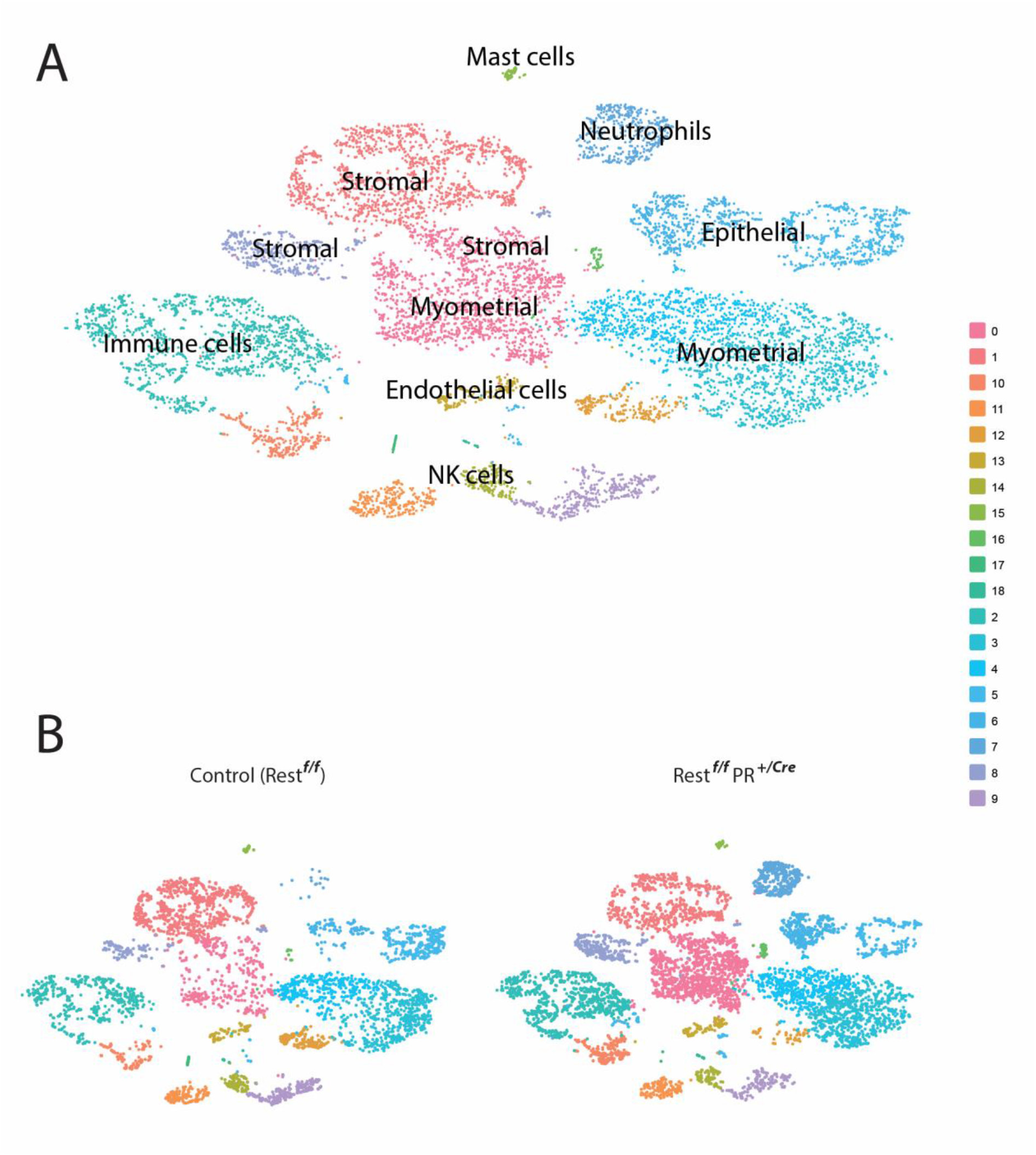
Single Cell RNA-sequencing analysis. (**A)** A t-SNE plot of the hierarchical clustering from uteri of 5-month-old control and *Rest*^f/f^ *PR^+/Cre^* cKO mice showed 19 distinct clusters. Populations of cells included epithelial, neutrophils, mast, immune, myometrial, endothelial, NK, and stromal cells. (**B)** Individual t-SNE plots of control and *Rest*^f/f^ *PR*^+/Cre^ cKO showed differences in subpopulations.

Uteri of *Rest^f/f^PR^+/Cre^* cKO mice showed increases in REST target gene expression, including overexpression of *Gria2* (Supplemental Figure 11A), *Stmn3* (Supplemental Figure 11B), and *Stmn2* (Supplemental Figure 11C). Additionally, upregulation of REST target genes was found in the stromal, epithelial, and myometrial cell clusters as represented in the violin plot (Supplemental Figure 12). Moreover, genes downstream of ERα signaling, including *Mmp24* (matrix metallopeptidase 24) and *Snap25* (synaptosome associated protein 25), were activated in the *Rest^f/f^PR^+/Cre^* cKO mice (Supplemental Figure 11D & 11E), indicating enhanced estrogen signaling upon the loss of REST. IPA analysis of stromal and myometrial cells (Cluster 0, fibroblast lineage) in cKO uteri showed gene expression profiles similar to those found in benign solid tumors (Supplemental Figure 13A). In addition, estrogen signaling genes were altered in this cluster (Supplemental Figure 13B). IPA also identified an adenomyosis phenotype in this cluster, which needs to be further explored (Supplemental Figure 13C). Lastly, the *Rest^f/f^PR^+cre^* cKO mice had higher levels of collagens and *Acta2* compared to control uteri (Supplemental Figures 14, 15). This phenotype is consistent with data from *Rest^f/f^ Amhr2^+/Cre^* cKO mice.

### Myometrial specific deletion of *Rest* Leads to Leiomyoma Phenotype

To faithfully recapitulate the loss of REST in human UL, we next developed a cKO mouse model in which REST was specifically deleted in the myometrial layer of the uterus. Using a proximal promoter sequence from the rat calbindin-D9K (*CaBP9K*) promoter, which drives transgene expression specifically in myometrium (8, 29), we generated myometrial specific Cre recombinase transgenic mice (CaBP9K-iCre, Myometrial specific M-iCre) (Fig 4). The Rest cKO mouse, *Rest^f/f^* M-iCre cKO, was generated by breeding the *Rest* ^f/+^ *MiCre* mice with the *Rest ^f/f^* mice.

Similar to the *Rest^f/f^ Amhr2^+/Cre^* mice, the *Rest*^f/f^ M-iCre cKO mice had increased uterine size, cystic glands, and abnormal uterine morphology compared to control uteri in diestrus (Figure 3E & F, Supplemental Figure 16). Additionally, we observed SMC tumors forming in the myometrial layer of the uterus of the *Rest*^f/f^ M-iCre cKO mice (Figure 3C & 4D). Furthermore, REST target genes *Stmn3, Stmn2*, and *Gria2* were all significantly overexpressed in the *Rest*^f/f^ M-iCre cKO mouse, confirming the loss of REST function (Figure 3G).

**3.**
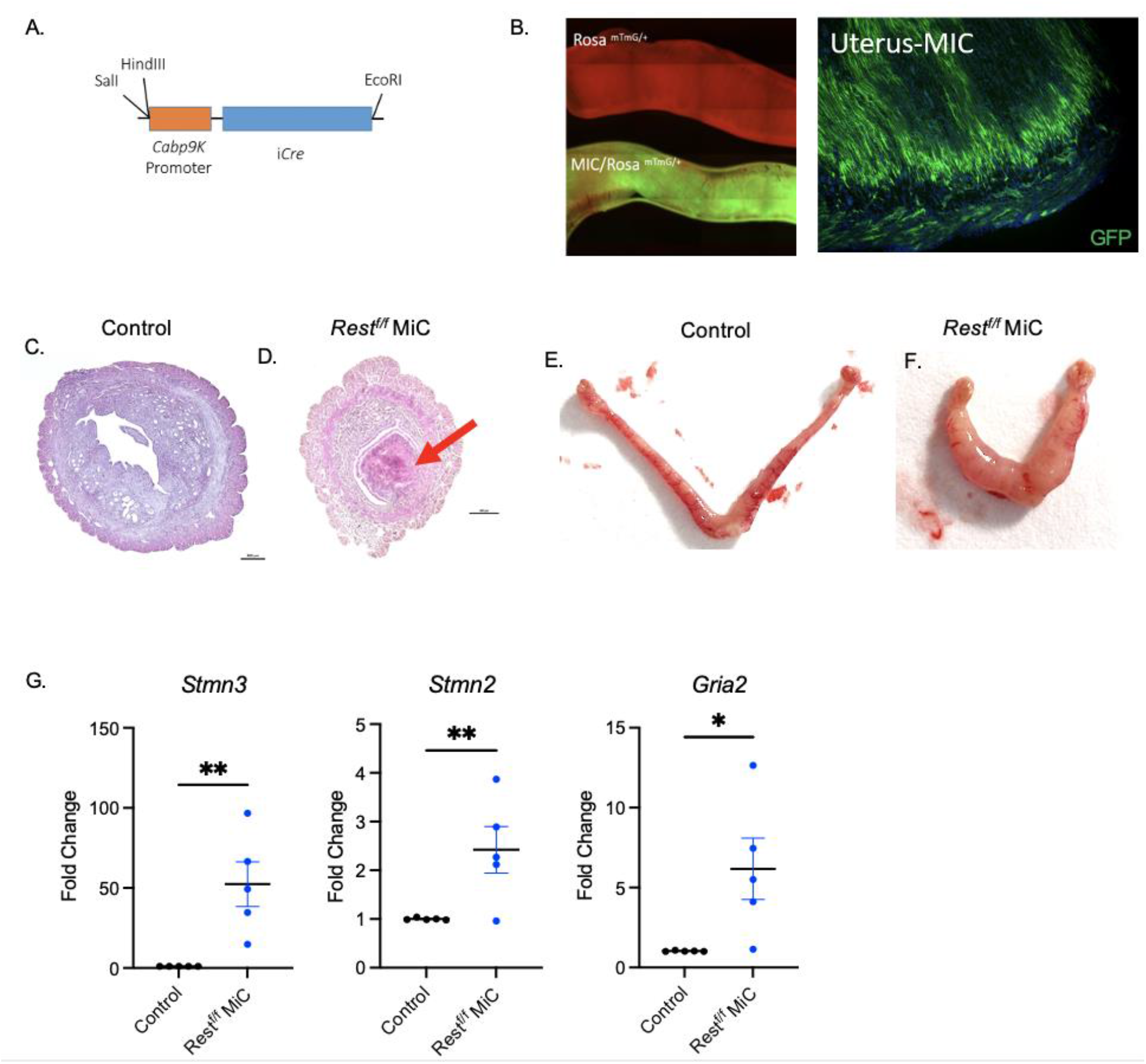
Generation and characterization of *Rest* cKO mouse under a myometrial specific (M-iCre) promoter. **(A)** Construct for the myometrial specific iCre expression using the proximal region of the rat *Cabp9k* promoter. **(B)** Left panel, top: Rosa^mTmG/+^ mouse uterine horn expressing tdTomato. Left panel, bottom: M-iCre/Rosa^mTmG/+^ mouse uterine horn expressing EGFP in the myometrium after tdTomato was removed using the myometrial specific iCre. Right panel shows EGFP expression is specific to the mouse myometrium. **(C)** H&E stain of uterine horn section in a control mouse. **(D)** H&E stain of uterine horn showing changes in morphology of *Rest^f/f^* M-iCre mouse. Tumor formation in the *Rest^f/f^* M-iCre mouse indicated by arrow. Magnification at 4x and scale bar 500um. **(E&F)** Representative image of increased uterus size of *Rest^f/f^* M-iCre mouse compared to control uterus during diestrus. **(G)** Gene expression of known Rest targets *Stmn3, Stmn2*, and *Gria2* in 6 month old *Rest^f/f^* M-iCre mice compared to control (n=5). Error bars represent +SEM. Student’s T-test was performed, *P<.05, **P<0.01

**Figure 4.**
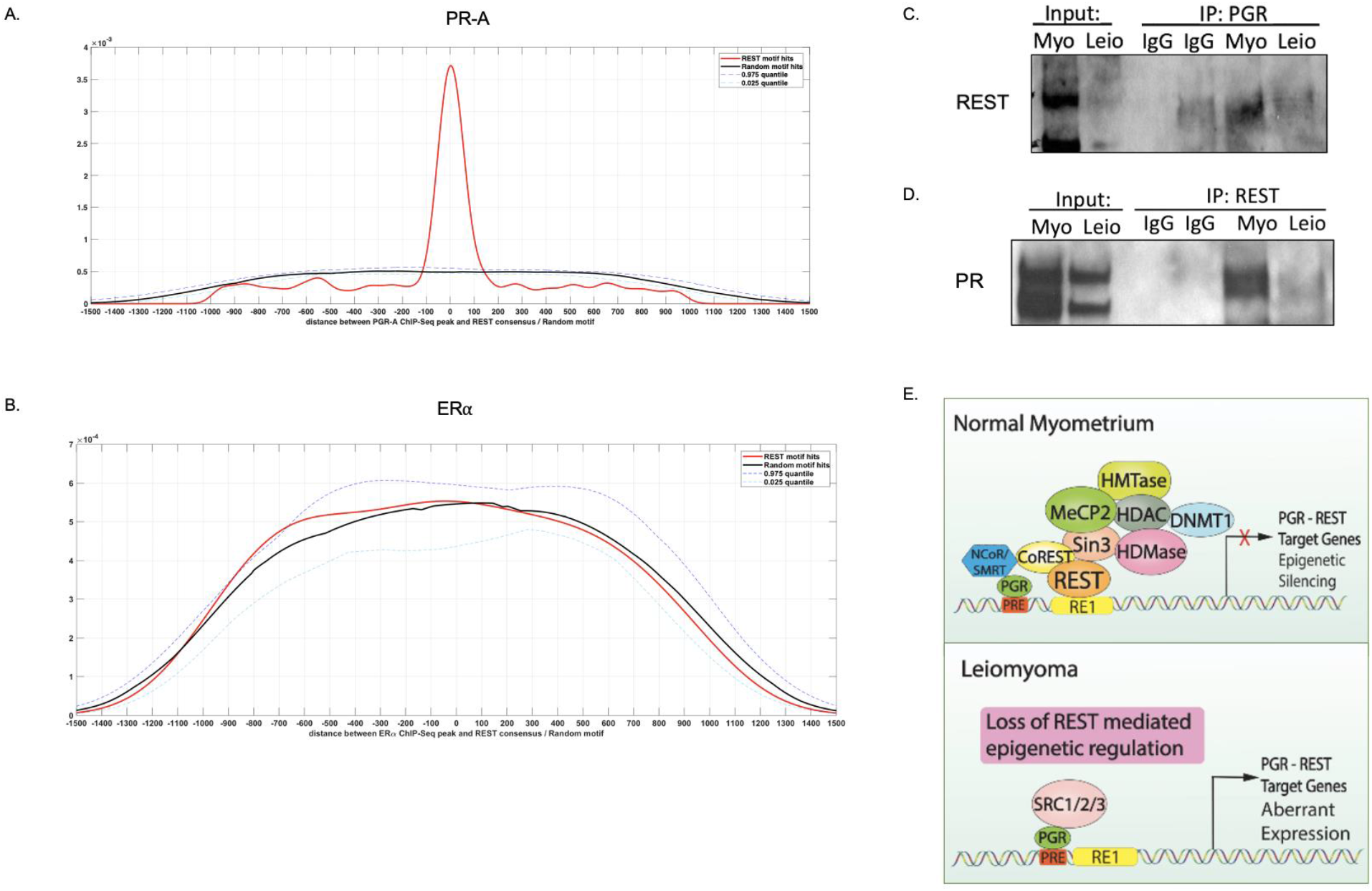
REST’s interactions with steroid hormone receptors. **(A)** A conserved frequency of occurrence of REST binding sites (0) within 100 base pairs of PR-A binding sites. X-axis represents distance in base pairs and y-axis represents density. **(B)** No overlap or proximal relationship between REST binding sites (0) and ERα binding sites. X-axis represents distance in base pairs and y-axis represents density. **(C&D)** Representative image of co-immunoprecipitation of REST and PR in human myometrial tissue and decreased interaction in leiomyoma tissue. **(E)** Working model depicting a REST-PR interaction in the normal myometrium and loss of REST’s impact in leiomyoma.

Uterine leiomyomas are known to be associated with altered expression of genes involved in TGF-β signaling, SMCs, and ECM components (30). Using uteri from 6-month-old mice, we tested the expression of *Tgfb3*, dermatopontin (*Dpt*), alpha smooth muscle actin (*Acta2*), and collagens *Col1A1* and *Col3A1* to determine if they contributed to the increased uterus size. Results indicated that the *Rest*^f/f^ M-iCre cKO mice expressed higher levels of *Col3A1*, and statistically significantly higher levels of *Dpt, Col1A1, Tgfb3*, and *Acta2* compared to control mice (Supplemental Figure 17). Dysregulation of these genes in the *Rest*^f/f^ M-iCre cKO mice, except *Dpt* that showed an increase in expression unlike the decrease reported in UL, represents a phenotype similar to human UL. Preliminary fertility data for the *Rest*^f/f^ M-iCre cKO female mice showed similar fertility compared to control mice (data not shown).

### REST Interacts with Progesterone Receptor

Our three *Rest* cKO models showed altered signaling, including activation of the estrogen receptor pathway as well as uterine hypertrophy during diestrus (Fig 1C) even when progesterone was present. To understand how REST regulated this phenotype, we investigated whether REST had a direct relationship with the steroid hormone receptors, ER and PR. Using available ChIP-Seq data sets for REST, PR-A, PR-B, and ERα (Supplemental Table 2, GSE62475, GSE36455), we identified a high frequency of conserved RE1 sites associated with REST within 100 base pairs of PR-A binding sites (Figure 4A) This conserved relationship was found on roughly 200 REST target genes. Additionally, analysis of binding sites of PR-B and REST in available ChIP-Seq data, identified conserved binding sites within 300 base pairs of each other (Supplemental Figure 18). Interestingly, there was no overlap or proximal relationship between ERα and RE1 binding sites (Figure 4B). These results indicate that the interaction between REST and PR is unique.

We next hypothesized that the absence of interaction between REST and PR due to the loss of REST could account for altered sex steroid hormone signaling seen in UL and in our mouse models. Results from co-immunoprecipitation (IP) studies with PR antibodies on healthy human myometrial tissue and paired UL specimen showed association of REST with PR in myometrial samples (Figure 4C). This interaction was reduced in the leiomyoma sample where REST expression was low (Figure 4C). Reciprocal immunoprecipitation with REST antibodies confirmed IP results with PR antibodies (Figure 4D). These results indicate that REST and PR interact in the healthy myometrium. However, upon the loss of REST, the protein-protein interactions between PR and its coregulators are altered, and lead to aberrant expression of PR-REST target genes (Figure 4E).

To further study the interaction between REST and PR, expression of REST-PR target genes that contained conserved binding sites were further investigated. Many of these targets were found to be dysregulated in the *Rest^f/f^ Amhr2^+/Cre^* cKO mice (Supplemental Table 4). Among the top 24 dysregulated REST-PR targets, which were also dysregulated in UL (Supplemental Table 5), the cell migration inducing hyaluronidase 1 (CEMIP, KIAA1199) contained both RE1 and PR binding sites within 1000 base pairs of each other (Figure 5A). We also found CEMIP to be significantly upregulated in human UL specimen compared to healthy myometrial specimen (Figure 5B) and correlated with low levels of REST (Figure 5B). Similarly, *Cemip* was found to be over expressed in the *Rest^f/f^ Amhr2^+/Cre^* cKO, and significantly overexpressed in the *Rest*^f/f^ M-iCre cKO mice uteri, at both the mRNA and protein levels (Figure 5C, D).

**Figure 5.**
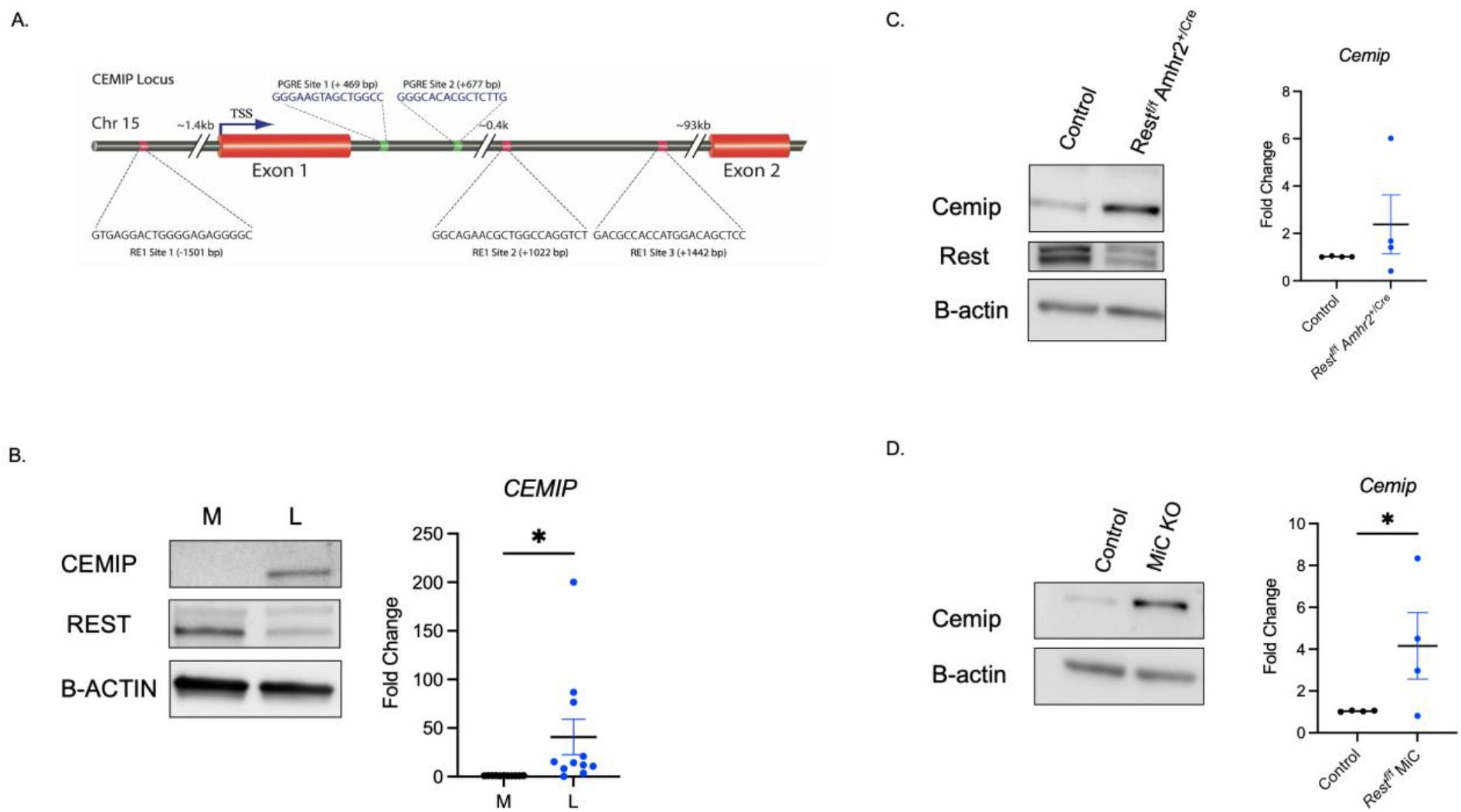
Overexpression of Cemip in uterine leiomyoma and *Rest* cKO mice. **(A)** CEMIP gene locus showing REST binding sites (RE1 sites) and PR binding sites (PGRE), and their locations relative to the transcription start site (TSS, +1). **(B)** Left panel: Western blot showing CEMIP overexpression and loss of REST in leiomyoma patient samples compared to control myometrium. Right panel: TaqMan RT-qPCR of *CEMIP* in leiomyoma patient samples compared to control myometrium (n=11). **(C) Left panel:** Western blot showing Cemip overexpression and loss of Rest in *Rest^f/f^ Amhr2^+/cre^* cKO compared to control. Right panel: TaqMan RT-qPCR of *Cemip* in *Rest^f/f^ Amhr2^+/cre^* cKO compared to control (n=4). **(D)** Left panel: Cemip overexpression shown by western blot in *Rest^fl/fl^* M-iCre cKO compared to control. Right panel: TaqMan RT-qPCR of *Cemip* in *Rest^f/f^* M-iCre cKO compared to control (n=4). B-actin was used as a loading control for western blots. Error bars represent +SEM. Student’s nonparametric T-test was performed, *P<0.05.

To investigate if REST directly affects expression of PR target genes, we focused on *CEMIP* expression, which is known to play a role in the regulation of hyaluronan (HA) and CD44 signaling (31, 32). Using siRNAs, REST was silenced in primary myometrial cells, and cells were treated with either vehicle or estradiol (10nM) and progesterone (1μm). After a 24-hour knockdown, *CEMIP* expression, at both gene and protein levels, was significantly increased in the siREST+ vehicle treated samples (Supplemental Figure 19A, B). In addition, CEMIP expression was altered in the absence of REST and presence of estrogen and progesterone compared to siControl-treated with hormones (Supplemental Figure 19B). While further studies are needed to determine the exact role CEMIP in UL pathogenesis, results of this experiment on one of the PR target genes showed how REST is involved in regulating target genes of PR. Further studies are needed to explore the molecular regulation of REST and PR on the expression of the *CEMIP* gene locus in the myometrium.

## Discussion

Despite the long-recognized roles of epigenetic and environmental factors that predispose women to UL, molecular events that lead to the initiation of UL growth and pathogenesis are still not well understood (33–35). Here we provide evidence that the loss of REST, a known tumor suppressor and a major epigenetic regulator of gene expression, plays a significant role in UL pathogenesis. We provide evidence that the loss of REST function leads to aberrant expression of REST target genes that contribute to cell proliferation and ECM accumulation in the uterus. We further provide evidence that loss of *Rest* in tissue-specific cKO mouse models leads to UL formation, and changes in uterine morphology and gene expression profiles with extensive similarities to human UL. Crucially, we show *Rest* cKO mice have altered responses to endogenous hormones, estrogen, and progesterone, due to a novel interaction between REST and PR.

REST is a master epigenetic silencer that represses neuronal genes in non-neuronal cells (36) through binding to a 21-23bp repressor element sequence (RE-1 site) located on an estimated 2,000 target genes within the human genome (37). Previously, we demonstrated that loss of REST leads to derepression of *GPR10 (PRLHR)*, which plays a role in UL pathogenesis by activating the PI3K/AKT-mTOR pathway (8, 38, 39). Additionally, *PRLHR* was shown to be overexpressed ubiquitously in UL specimen carrying wide range of known mutations or translocations (4), indicating that loss of REST may be a crucial upstream event in uterine fibroids (8). Our data demonstrate that loss of *Rest in vivo* leads to tumorigenesis in uterine SMCs and accumulation of ECM.

Preclinical models that accurately recapitulate altered steroid hormone pathways in UL, which are necessary for the development of safe and efficacious treatments for UL, are nonexistent in the field. Although Eker rats, which harbor a mutation in the *Tsc2* gene (40, 41) develop spontaneous, estrogen-sensitive UL tumors, they also develop fatal renal and liver cancers (42). Additionally, UL growth in this model can only be induced by estrogen and not progesterone (43). In addition, formation of UL in another model in guinea pig is also dependent on estrogen (44). These tumors do not have similar histological features to human UL and are inhibited by progesterone treatment (3). Although recently developed xenograft model of UL shows tumor growth in the presence of estrogen and progesterone (45), this model is not useful to study initiation and development of UL. There is, therefore, an urgent need for UL-relevant animal models which are sensitive to estrogen and progesterone, and represent cellular, molecular, and genetic features of human UL; the *Rest*^f/f^ M-iCre cKO and *Rest^f/f^ Amhr2^+/Cre^* cKO mouse models fulfil this unmet need in the field.

Specifically, the *Rest* cKO mice showed increased uterine size, SMC proliferation and ECM deposition, which are key features of human UL tumors. Similar to increased sensitivity of human UL to estrogen and the abnormal expression of estrogen responsive genes during the luteal phase (1), the cKO mice showed increases in uterine size during diestrus when estrogen and progesterone were both present. Moreover, results from the single cell RNA sequencing of the *Rest^f/f^ PR^+/Cre^* cKO mice provided compelling evidence that loss of REST has significant effects on uterine tissue architecture. While Amhr2^+/Cre^ and PR^+/Cre^ drivers are traditionally used to delete floxed genes in the reproductive tract, we developed and utilized a novel myometrial specific Cre driver to delete *Rest* in the myometrium, to further confirm a distinct role of REST in UL pathogenesis. This Cre driver (CaBP9K-iCre, M-iCre) will be invaluable for future studies involving myometrial specific gene deletion.

Importantly, our data show that the loss of REST can lead to alterations in sex steroid hormone signaling, and UL development. Our results showed (a) REST and PR interact in the heathy myometrium, and (b) REST and PR target genes are dysregulated in UL. One of the novel targets of REST, CEMIP, is significantly overexpressed in both human UL and in our cKO mouse models. CEMIP has been shown to induce fibrosis in arthrofibrosis (46), and its upregulation in several cancer types is linked to cell proliferation, migration and changes in cell signaling, such as PI3K/AKT (47, 48), which is overactivated in UL. In addition, CEMIP is a HA binding protein involved in HA depolymerization (31). HA is ubiquitously present in the ECM and provides structural integrity to the uterus. It has also been shown that under certain pathological conditions HA degradation is enhanced, and the lower molecular weight molecules, resulting from HA degradation, can contribute to tumor growth and angiogenesis (49). We recently showed preferential association of REST with RE1 sites within the CEMIP locus as well as aberrant expression of CEMIP in the absence of REST in breast cancer cells (50). Based on our results showing loss of REST leads to overexpression of CEMIP, it will be important to test what role CEMIP plays in UL pathogenesis and how PR and progesterone affect this gene.

Current hormone therapies for UL suffer from poor safety profiles, precluding them from long-term use (51, 52). Current generation of SPRMs, which are developed based on PGR antagonism, reduce UL tumor size in randomized control trials (16, 53). However, side effects of endometrial hyperplasia have been reported and concerns of their effects on non-targeted tissues (breast, ovary, liver) exist (53). Our work provides a novel link between the altered response to progesterone and loss of REST in UL. We have found that REST interacts with PR and influences the expression of target genes. We believe this unique relationship between REST and PR presents a mechanism that can be targeted to develop a new generation of SPRMs.

## Methods

### Study approval

All human studies were approved by and performed in accordance with the University of Kansas Medical Center Policies and Procedures Relating to Human Subjects (IRB#: 5929). All patients gave informed consent when donating their tissue samples to the University of Kansas (KU) Cancer Center’s Biospecimen Repository Core Facility. All mouse experiments were approved by the University of Kansas Medical Center IACUC protocols (2019-2508) and adhere to NIH guidelines for care and use of laboratory animals.

### Tissue collection and cell culture

Matched myometrium and leiomyoma tissue samples were obtained from pre-menopausal women undergoing hysterectomies at the University of Kansas Hospital (Kansas City, KS). Criteria for patients in this study exclude women undergoing hysterectomy for a primary condition other than uterine leiomyoma and those taking hormone therapy in the three months preceding surgery. Myometrial primary cells were prepared from samples as described previously (8).

### Generation of mouse models, gene expression analysis

Details of the generation of mouse models, gene and protein expression analysis are described in “Supplemental Methods” section.

## Author Contributions

VMC designed research; FK, MMM, ASC, SGa, SGu, AG performed research; FK, MMM, ASC, SGu, VMC analyzed data; SGu, WBN contributed reagents/materials; and ASC, VMC wrote the manuscript. FK, WBN, SGu made edits to the manuscript. ASC, FK, MMM contributed equally to the manuscript.

## Acknowledgments

VMC was supported by grants from the NIH: P20 RR016475, R01 HD094373, R01HD076450. Authors acknowledge University of Kansas (KU) Cancer Center Biospecimen Repository Core for human specimens, KU Cancer Center’s Support Grant (P30 CA168524), Genomics Core supported by the Kansas Intellectual and Developmental Disability Research Center (NIH U54 HD090216), COBRE (P30 GM122731-03) and the NIH S10 High-End Instrumentation Grant (NIH S10 OD021743) at KUMC, Kansas City, KS 66160. We thank The University of Virginia Center for Research in Reproduction Ligand Assay and Analysis Core, supported by the NICHD/NIH (NCTRI) Grant P50-HD28934.

## Notes

**Conflict of interest**, The authors have declared that no conflict of interest exists.

### Competing Interest Statement

The authors have declared no competing interest.

