## Supplemental file for "Loss of the repressor REST affects progesterone receptor function and promotes uterine leiomyoma pathogenesis"

**Supplemental Methods**

Generation of *Rest* conditional knockout mice

Floxed *Rest* embryonic stem cell clones were obtained from the European Conditional Mouse Mutagenesis Program (http://www.eucommm.org). The cells were used to generate chimeric founder mice, which were mated to C57BL6 wild type mice to create *Rest* fl/+ mice. FLPe expressing mice (B6.129S4-Gt(ROSA) 26Sor$^{tm1(FLP)Dym}$/RainJ) (JAX stock #009086) were then crossed with *Rest*$^{f/+}$ mice to remove FRT flanked βgal-neo sequences (1). Restfl/fl mice were obtained and crossed with *Amhr2*$^{+/Cre}$ recombinase mice (2) or with PR$^{+/Cre}$ mice (3) to produce *Rest* conditional knockout (cKO) mice in the female reproductive tract. The mice were genotyped by PCR using forward *Rest* primer (5'TGTAGTTTCCAAACTGTGACTTCG) and two reverse primers (5'TGAACTGATGGCGAGCTCAGACC) (5'GCTACAAAATGCGTAAGTTCAAGG), in addition to primers for *Amhr2* (5'GGACATGTTCAGGGATCGCCAGGC) (5'CGACGATGAAGCATGTTTAGCTG).

Myometrial specific iCre (M-iCre) generation

A pGEMT vector containing iCre with a sv40 polyA tail was obtained from Dr. Rajendra Kumar (4). The rat calbindin D9K short (-117 to +365bp) promoter (Genebank: X16635.1) was PCR amplified from our previous vector (5) with primers containing a AatII and NotI site at the 5' end and a reverse primer containing a SacII site. The following primers were used for gene amplification/cloning: forward primer (Cabp9k): AGTCGACGTCGCGGCCGCGCTCAAGCTTGGTCTCAGA and reverse primer ATGCCCGCGGTTTTCTGTGCTGTAAGGGAAG.

The amplified promoter sequence was digested using AatII and SacII restriction enzymes. The digested insert was then ligated into the iCre -pGEMT vector using T4 DNA ligase. The PCR product was cloned into the vector downstream of a NotI restriction site. After sequencing to confirm the integrity of the construct, NotI was used to release the insert. The insert containing the promoter, iCre and sv40 polyA after gel elution and purification was used to generate the myometrial specific iCre (M-iCre) expressing mice. M-iCre mice were mated with *Rest*$^{f/f}$ mice and F1 progeny were born. Mice were genotyped by PCR using M-iCre forward primer (5’CCACTAATGCTGTTCCGACCTGTC) and reverse primer (5’CATCCTTGGCACCATAGATCAG) and forward *Rest* primer (5’TGTAGTTTCCAAACTGTGACTTCG) and two reverse primers (5’TGAACTGATGGCGAGCTCAGACC) (5’GCTACAAAATGCGTAAGTTCAAGG).

Histology and staining

Uterine tissues were fixed in 4% paraformaldehyde and processed for paraffin embedding. Tissue sections were deparaffinized in xylene, rehydrated through a series of ethanol and stained with Hematoxylin and Eosin (H&E). For immunofluorescence, rehydrated tissue sections were subjected to staining as described previously (5), using primary antibodies COL3A1 (Novus Biologicals, #NB600-594SS) and αSMA (Millipore, #MABT381).

RNA isolation and qRT-PCR analyses

Total RNA was isolated from tissue samples or cultured cells stored in RNAlater (Qiagen, Valencia, CA) using RNeasy Mini Kit (Qiagen, Valencia, CA) according to manufacturer’s protocol. After quantification using Nanodrop spectrophotometer, aliquots of RNA were reverse transcribed using High-Capacity cDNA Reverse Transcription Kit (Applied Biosystems, Waltham, MA). TaqMan assays for *Rest* (IDT, Mm.PT.58.12166480), *Gria2* (IDT, Mm.PT.58.8802331), *Stmn3* (IDT, Mm.PT.58.23492094), *Stmn2* (IDT, Mm.PT.58.13787385), *Col1A1*(IDT, Mm.PT.47.6999992), *Dpt* (IDT, Mm.PT.47.17098032), *Tgfb3* (IDT, Mm.PT.47.10648587), *Col3A1* (IDT, Mm.PT.47.9778198), *Acta2* (IDT, Mm.PT.47.7024949), *Cemip* (ThermoFisher Scientific, Mm00472921_m1), and *CEMIP* (IDT, Hs.PT.58.28305095) were used to quantify gene expression differences utilizing the delta delta C(T) method with housekeeping genes Rn18s (ThermoFisher Scientific, Mm03928990_G1) or 18s (IDT, Hs.PT.39a.22214856.g).

Protein extraction, western blotting, and co-immunoprecipitation

Frozen tissue samples were bio pulverized. Tissue samples were then lysed using 2x cell lysis buffer (Cell Signaling Technology, Danvers, MA) supplemented with protease and phosphatase inhibitor cocktails (ThermoFisher Scientific, Waltham, MA). Cultured cells were lysed using 1x cell lysis buffer (Cell Signaling Technology, Danvers, MA) supplemented with protease and phosphatase inhibitor cocktails (ThermoFisher Scientific, Waltham, MA). Samples were then sonicated and centrifuged at 18,000g at 4°C. Protein was quantified using Bio-Rad Protein Assay (Bio-Rad, Hercules, CA). Western blots were performed as described previously (5). Proteins were detected using SuperSignal West Pico PLUS Chemiluminescent Substrate (ThermoFisher Scientific, Waltham, MA) and blots were imaged using BioRad ChemiDoc MP (BioRad, Hercules, CA). The following antibodies were used: REST (Millipore, #07-579), GPR10 (Novus Biologicals, NBP1-00854), AKT (Cell Signaling Technology, #9272), P-AKT (Cell Signaling Technology, #40605), P-4EBP1 (Cell Signaling Technology, #9459s), P-P70S6K (Cell Signaling Technology, #4206s), GAPDH (Cell Signaling Technology, #5174s), PR (Cell Signaling Technology, #8757s), Beta-actin (Genescript, #A01865), CEMIP (Novus Biologicals, #45750002) and anti-rabbit-HRP (Promega). Co-immunoprecipitations were done using Pierce Crosslink Magnetic IP/Co-IP Kit (ThermoFisher Scientific, Waltham, MA) according to manufacturer’s protocol. Antibodies to REST (Millipore, #17-641), PR (ThermoFisher Scientific, #MA1-12626), and Rabbit IgG (Cell Signaling Technology, negative control) were used.

Hormone assessment

Samples of serum from mice at diestrus were assayed by ELISA at the University of Virginia Center for Research in Reproduction, Ligand Assay and Analysis Core (Charlottesville, VA).

Transfections

Cultured primary myometrial cells were plated in 6-well plates at 300,000 cells per well in antibiotic free media. After 24 hours at 37$^{o}$C, 5%$CO_2$, cells were transfected with a 100pmol mixture of Silencer Select siRNAs (S11934, S11932, S119323) to *REST* from Ambion (Life Technologies, Carlsbad, CA) using Lipofectamine 2000 transfection reagent (Invitrogen, Carlsbad, CA) according to manufacturer's protocol. Control experiments included ON-TARGETplus nontargeting scrambled siRNA 2 (ThermoFisher Scientific, Waltham, MA). After 24 hours, transfected cells were treated with vehicle (PBS) or estradiol (10nM) and progesterone (1uM) in charcoal stripped serum, phenol red free medium (Lonza, Morristown, NJ). After 24 hours of treatment, RNA and protein extracts were collected and analyzed.

RNA sequencing

RNA isolation from whole uteri of *Rest*$^{f/f}$ *Amhr2*$^{+/Cre}$ cKO and control mice aged 4 months was performed using phenol: chloroform. RNA samples were prepared by the KUMC Genomics Core using the TruSeq Stranded Total RNA LT sample Preparation Kit with Ribo-Zero Mouse. Libraries were adjusted using the Agilent 2100 Bioanalyzer using the Agilent DNA 1000 Chip. Sequencing was performed on the Illumina HiSeq2500 sequencing System (Illumina, Inc., San Diego, CA). Data deposited in the NCBI Gene Expression Omnibus (GEO, GSE178141). The samples were analyzed in biological triplicates giving six samples in total. Each sample generated between 109.5 and 128.1 million fragments (reads). Reads were mapped to the mouse genome (GRCm38) using the STAR software (6) (version 2.3.1z). Between 80.4% and 88.4% of the sequenced reads mapped to the mouse genome. Differential gene expression analysis was performed using Cuffdiff (7) (version 2.1.1 2). The resulting p-values were adjusted for multiple hypothesis testing using the Benjamini and Hochberg method (8).

Single cell RNA sequencing

Whole uteri of *Rest*$^{f/f}$ *PR*$^{+/Cre}$ cKO and control mice aged 5 months were isolated immediately after euthanasia using IACUCC approved methods and washed in cold 1X PBS. Uteri were processed by mincing the tissue and digesting in DMEM (Gibco, Waltham, MA), 10% FBS (Atlanta Biologicals, Flower Branch, GA), 1.5mg/ml collagenase (Gibco, Waltham, MA), and 2400U of DNaseI (NEB, Ipswich, MA) for 5 hours in a shaking water bath at 37°C. After digestion, cells were centrifuged at 300g for 5 minutes and treated with ACK lysis buffer followed by two washes with 1X PBS and 2% FBS. Cells were passed through a 100um cell strainer and resuspended in 1X PBS and 2% FBS. Cells were passed through a 40um cell strainer and resuspended in 1X PBS and 2% FBS. Control samples (n=2) were pooled and cKO samples (n=2) were pooled. The KUMC genomics core determined concentration and viability using the Countess II FL Automated Cell Counter (Invitrogen, Waltham, MA) followed by single cell library preparation using the 10x Chromium Controller (10X Genomics, Pleasanton, CA). Sequencing was done using the NovaSeq 6000 (Illumina, Inc., San Diego, CA). The transcriptomic profiles of Restf/f PR+/Cre cKO and control mice at a single-cell resolution were obtained using the 10x Genomics Chromium Single Cell Gene Expression Solution (10xgenomics.com). The primary analysis of the scRNAseq data was performed using the 10x Genomics Cell Ranger pipeline (version 5.0.1). This pipeline performs sample de-multiplexing, barcode processing, and single cell 3’ gene counting. The cKO sample consisted of 9,075 cells with 50,201 (mean) reads and 1,275 (median) genes per cell. The control sample consisted of 6,082 cells with 85,407 (mean) reads and 1,804 (median) genes per cell. Both samples achieved high sequence saturation levels. The mapping rate of reads to the mouse genome (mm10) was around 93% for both samples. Cells identified as doublets using the DoubletFinder software (9) were removed prior to analysis. Cells were further filtered on the number of UMIs (> 500), number of genes detected (> 250 and < 5000), number of genes per UMI ($\log_{10}$ value > 0.8) and mitochondrial transcript ratio (< 0.2). The single-cell data was analyzed using the Seurat[1] R package. The SCTransform (10) method in the Seurat package was used for data normalization and variance stabilization. The Seurat software identified 19 cell clusters that were visualized via t-distributed stochastic neighbor embedding (t-SNE). The cluster cell types were identified using the SingleR (11) package in R and expert curation based on marker genes. The data were visualized and curated with the Loupe Browser software (10X Genomics, Pleasanton, CA) after uploading the analysis information from the Seurat package. Stromal cells were negative for *Epcam* (epithelial cell adhesion molecule) and *Cd45* (protein tyrosine phosphatase receptor type C) but positive for *Ifitm1* (interferon induced transmembrane protein 1), *Lum* (Lumican), *Vit* (Vitrin), *Ngfr* (nerve growth factor receptor), and *Pdgfrb* (platelet derived growth factor receptor beta). Epithelial markers included *Krt18* (keratin 18), *Krt19* (keratin 19), and *Epcam*. Myometrial cells showed high levels of *Acta2* (actin alpha 2, smooth muscle), *Cnn1* (calponin 1), *Cald1* (caldesmon 1), *Tpm1* (tropomyosin 1) and *Vim* (vimentin). Immune, mast, NK, and neutrophil cell types were predicted using the immunological genome project (ImmGen) and SingleR. The FindMarkers function in the Seurat package with default parameters were used for differential expression analysis. This data is deposited in GEO (GSE178141).

Distribution of REST binding elements in the vicinity of PGR-A, ERα and PGR-B binding regions

PGR/ER ChIP-Seq binding regions identified by MACS (12) were centered at the location of significant enrichment of ChIP-Seq tags (peak summit). The genomic region extending 1000 bases on either direction from the summit was obtained. REST binding elements within this region were identified using a sequence specific weight matrix for the REST motif (> 80% identity). The distances from these elements to the PGR peak summit were recorded. The empirical probability density estimate of these distances were calculated using a kernel density estimator with a normal kernel function. The empirical background distribution was established by repetitively performing the above procedure for 100 random motifs with the same nucleotide composition as the REST motif. The density estimates were plotted where the solid red graph represents the estimated empirical probability density of the distance of REST binding elements from the PGR/ER ChIP-Seq peak. The solid black graph represents the estimated empirical probability density of the median distance of random binding elements from the PGR/ER ChIP-Seq peak. The perforated blue lines represent the 0.975 and 0.025 quantiles of the estimated empirical probability density of the random motif distances to the peak.

Data mining and ingenuity pathway analysis

GEO dataset GSE13319 (13) was analyzed for the expression of REST associated / regulated genes in myometrial and leiomyoma samples. The dataset was background corrected, normalized and gene-level summarized using the Robust Multichip Averages Procedure (RMA). Statistical analysis was performed on biological triplicates. Biological, functional and pathway analysis were performed using Ingenuity Systems Pathway Analysis software (IPA, QIAGEN, Germantown, MD) on the significantly (FC $\geq$1.5, p-value $\leq$0.05) differentially expressed genes between myometrium and leiomyoma, or between wild type and cKO mouse. In addition, ChIP-seq datasets (Supplemental Table 2) from ENCODE (14) were used to determine potential REST target genes by examining REST binding to RE1 sites located within or near genes of interest. REST DNA binding sites were identified by the presence of both a strong ChIP-peak and REST consensus sequence within the peak region with at least 80% identity. GSE62475, GSE48096, GSE36455 were used to determine potential PR, ERβ, and ERα target genes respectively by examining PRE and ERE sites located within or near genes of interest. DNA binding sites were identified by the presences of both a strong

ChIP-peak and PRE or ERE consensus sequence within the peak region with at least 80% identity.

### Statistical analysis

Leiomyoma samples were compared to myometrial controls from the same patient. For animal studies, *Rest* cKO mice uterine samples were compared to age-matched control mice at the same stage of the estrous cycle. Quantitative experiments were repeated with at least three independent biological replicates. Statistical significance was determine using students T-test to determine change from the control sample. For paired tissue samples, the study was powered to measure changes in gene expression between the tumor and normal tissues. Using paired t-test, 11 pairs gave us 80% power to detect a difference of 0.81 (FC of 1.75) standard deviation units or larger in gene expression with one-sided level of significance of 0.05. Significance was set at $P<0.05$ for all comparisons. For RNA- sequencing results, genes used in IPA were filtered for false discovery rate (FDR) <0.05 and fold change of 1.5 or higher. Concordance between RNA- Sequencing results of $Rest^{f/f}$ $Amhr2^{+/Cre}$ cKO mice and human microarray data were analyzed by the hypergeometric statistical analysis test of the significance of the number of genes significantly differentially expressed in the same direction in both experiments (p-value < 3.24E-14).

### Supplemental Figures

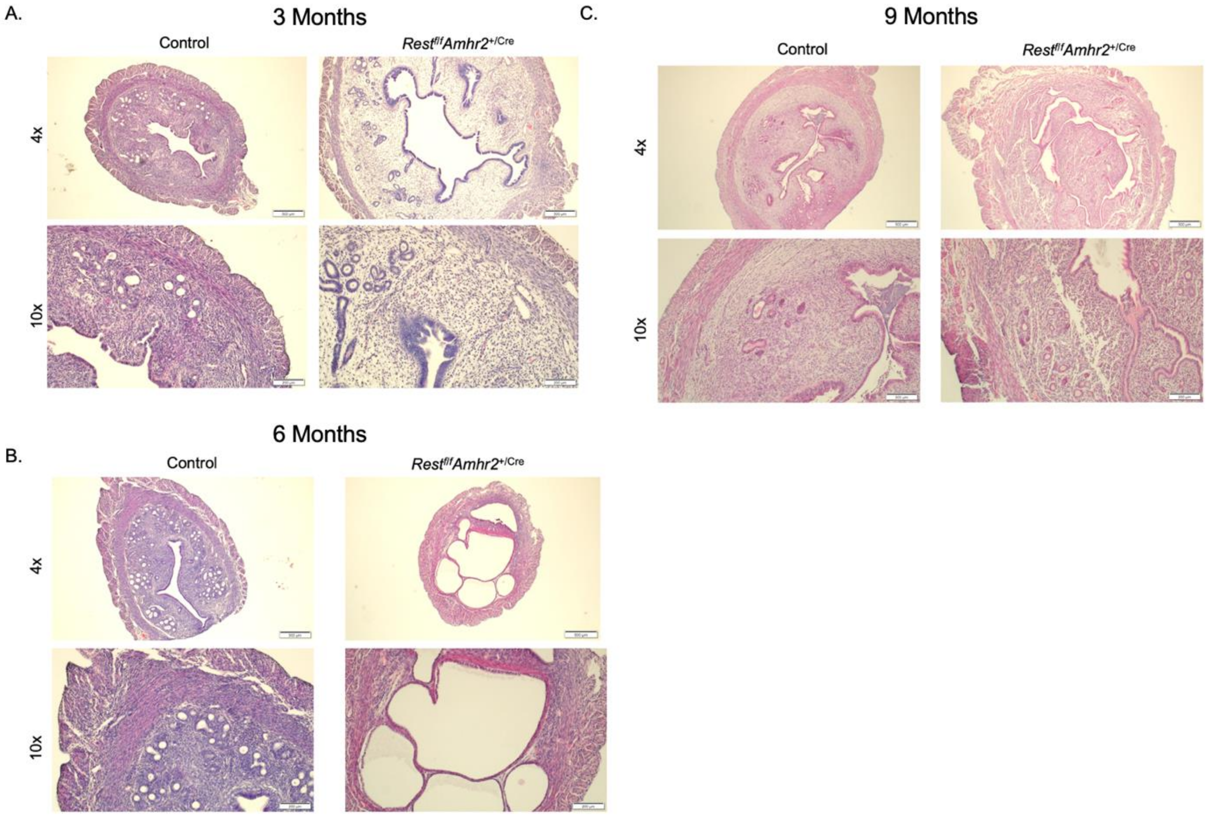


**Supplemental Figure 1. Characterization of uteri in *Rest*$^{f/f}$ *Amhr2*$^{+/Cre}$ mice.**
Uterus of control and *Rest*$^{f/f}$ *Amhr2*$^{+/Cre}$ mice at (A) 3, (B) 6, and (C) 9 months, in similar stages of the estrous cycle

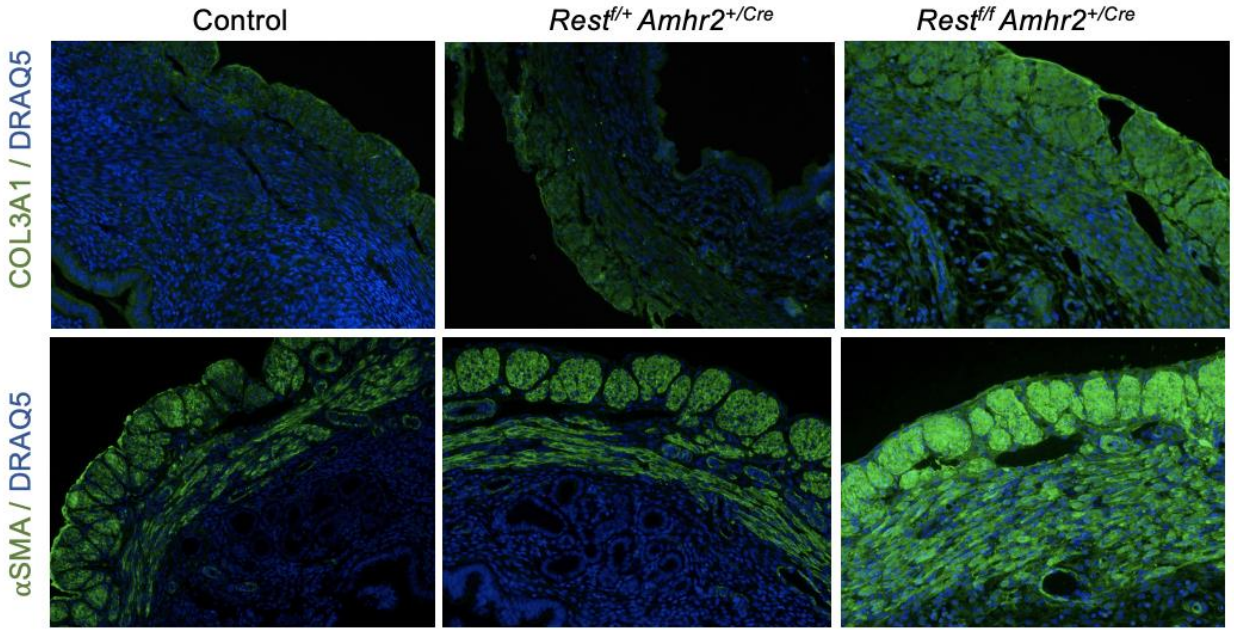


**Supplemental Figure 2. Characterization of uterine myometrium in control, *Rest$^{f/+}$ Amhr2$^{+/Cre}$*, and *Rest$^{f/f}$ Amhr2$^{+/Cre}$* cKO mice.** Green: Expression of Col3A1 and alpha-smooth muscle actin (αSMA); Blue: Nuclear staining with DRAQ5 (10x magnification)

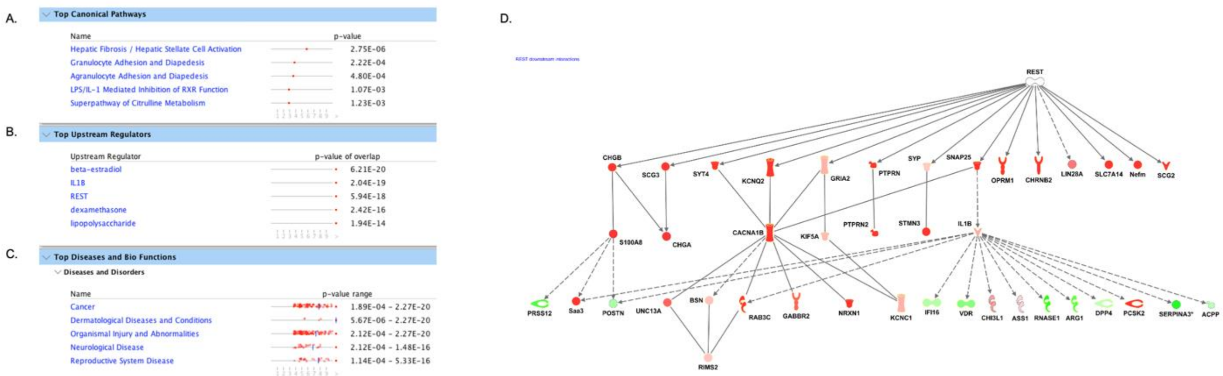


**Supplemental Figure 3. Altered genes, regulators and pathways in *Rest$^{f/f}$ Amhr2$^{+/Cre}$* cKO uteri. (A)** Top canonical pathways affected in the *Rest$^{f/f}$ Amhr2$^{+/Cre}$* cKO mice as predicted by the IPA, which is based on genes with a significant (P <0.05) differential expression compared to control mice. **(B)** Top upstream regulators of Rest predicted by the IPA to be dysregulated in *Rest$^{f/f}$ Amhr2$^{+/Cre}$* cKO mice compared to control mice **(C)** Top diseases and functions impacted in *Rest$^{f/f}$ Amhr2$^{+/Cre}$* cKO mice compared to control mice, per the IPA. **(D)** IPA network analysis on direct and indirect REST-associated genes in *Rest$^{f/f}$ Amhr2$^{+/Cre}$* cKO mice predicted to besignificantly (p <0.05) upregulated (red) or down regulated (green) in the whole uteri of 4-month-old *Rest$^{f/f}$ Amhr2$^{+/Cre}$* cKO mice in diestrus compared to control (n=3).

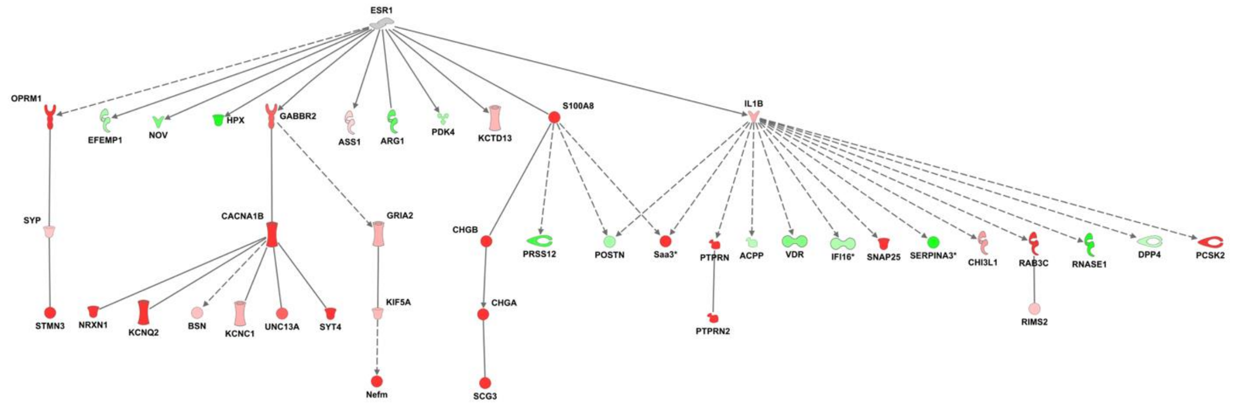


**Supplemental Figure 4. Altered hormone signaling in *Rest*$^{f/f}$ *Amhr2*$^{+/Cre}$ cKO uteri.** Gene network analysis of ESR1-associated genes per the IPA in GSE178141 (whole uteri genes during diestrus) predicting significantly ($p < 0.05$) upregulated (red) or down regulated (green) genes in 4-month-old *Rest*$^{f/f}$ *Amhr2*$^{+/Cre}$ cKO mice compared to control (n=3).

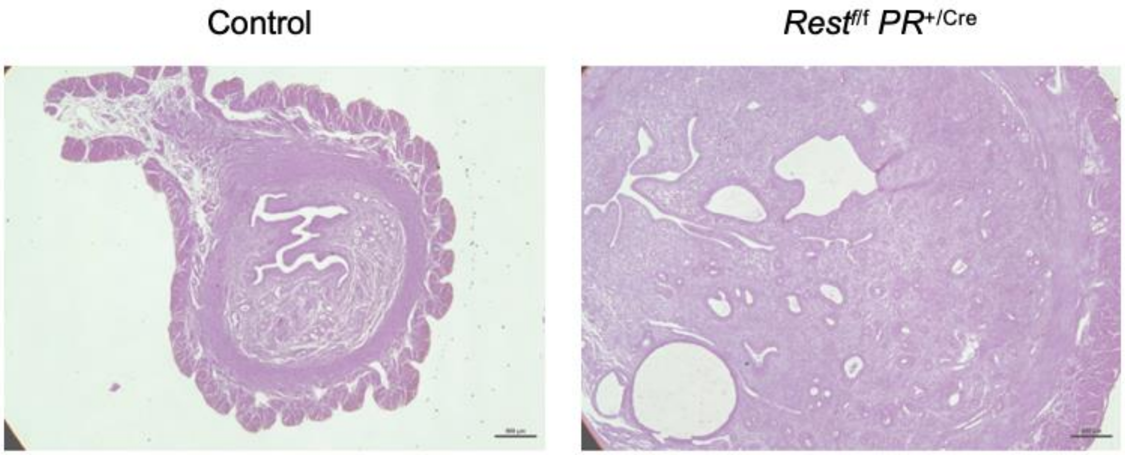


**Supplemental Figure 5. Characterization of *Rest$^{f/f}$ PR$^{+/Cre}$* uteri.** Representative images of H&E staining of uterine horn sections in *Rest$^{f/f}$ PR*$^{+/Cre}$ mouse compared to control. Scale bar: 500µm. (4x magnification)

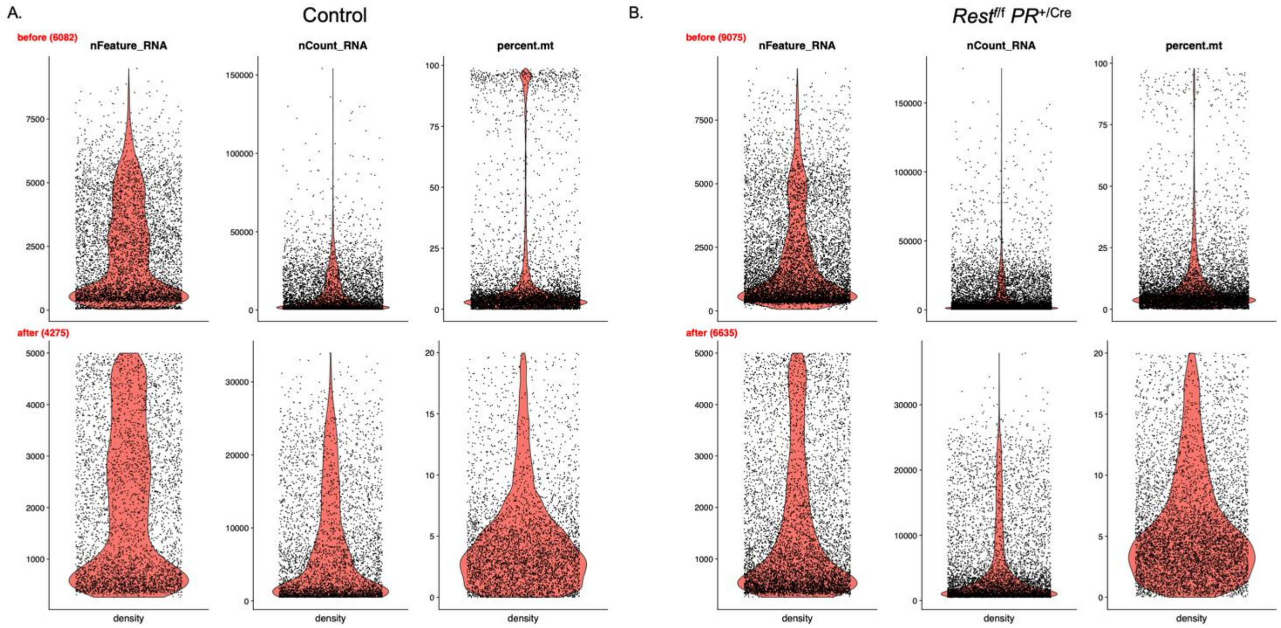


**Supplemental Figure 6. Quality control distributions for single cell RNA sequencing.** Violin plots depicting the number of features (nFeature_RNA), number of unique molecular identifiers (nCount_RNA) and percentage reads mapped to mitochondrial genes (percent.mt) in each cell, before and after QC filtering, in **(A)** Control and **(B)** *Rest*$^{f/f}$ *PR*$^{+/Cre}$ uterine samples.

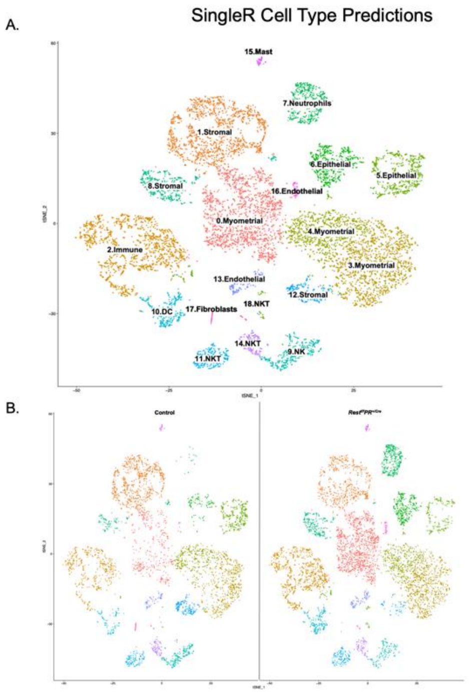


C.

| Cluster | Cluster SingleR (main) | Sample | Number of cells | Expected number of cells | Percentage deviation | Bonferroni pvalue |
|---|---|---|---|---|---|---|
| 0 | Fibroblasts | Control | 384 | 679 | -43.451 | 0 |
| 0 | Fibroblasts | Restf/f PR+/Cre | 1349 | 1054 | 27.996 | 0 |
| 1 | Stromal | Control | 831 | 574 | 44.761 | 0 |
| 1 | Stromal | Restf/f PR+/Cre | 634 | 891 | -28.840 | 0 |
| 2 | Monocytes | Control | 459 | 562 | -18.370 | 7.7865E-08 |
| 2 | Monocytes | Restf/f PR+/Cre | 976 | 873 | 11.836 | 7.7865E-08 |
| 3 | Myometrial | Control | 571 | 543 | 5.138 | 1 |
| 3 | Myometrial | Restf/f PR+/Cre | 815 | 843 | -3.311 | 1 |
| 4 | Myometrial | Control | 394 | 358 | 10.132 | 0.38957609 |
| 4 | Myometrial | Restf/f PR+/Cre | 519 | 555 | -6.528 | 0.38957609 |
| 5 | Epithelial | Control | 382 | 272 | 40.473 | 0 |
| 5 | Epithelial | Restf/f PR+/Cre | 312 | 422 | -26.077 | 0 |
| 6 | Epithelial | Control | 118 | 205 | -42.310 | 7.17204E-14 |
| 6 | Epithelial | Restf/f PR+/Cre | 404 | 317 | 27.261 | 7.17204E-14 |
| 7 | Neutrophils | Control | 16 | 180 | -91.104 | 0 |
| 7 | Neutrophils | Restf/f PR+/Cre | 443 | 279 | 58.699 | 0 |
| 8 | Stromal | Control | 119 | 165 | -27.864 | 0.000108853 |
| 8 | Stromal | Restf/f PR+/Cre | 302 | 256 | 17.953 | 0.000108853 |
| 9 | NK cells | Control | 247 | 156 | 58.780 | 0 |
| 9 | NK cells | Restf/f PR+/Cre | 150 | 241 | -37.872 | 0 |
| 10 | B Cells | Control | 111 | 137 | -18.832 | 0.155884954 |
| 10 | B Cells | Restf/f PR+/Cre | 238 | 212 | 12.133 | 0.155884954 |
| 11 | NKT | Control | 137 | 115 | 18.922 | 0.314962783 |
| 11 | NKT | Restf/f PR+/Cre | 157 | 179 | -12.192 | 0.314962783 |
| 12 | Myometrial | Control | 208 | 101 | 106.547 | 0 |
| 12 | Myometrial | Restf/f PR+/Cre | 49 | 156 | -68.649 | 0 |
| 13 | Endothelial cells | Control | 84 | 75 | 11.652 | 1 |
| 13 | Endothelial cells | Restf/f PR+/Cre | 108 | 117 | -7.508 | 1 |
| 14 | NKT | Control | 103 | 74 | 38.348 | 0.000709062 |
| 14 | NKT | Restf/f PR+/Cre | 87 | 116 | -24.708 | 0.000709062 |
| 15 | Mast cells | Control | 27 | 23 | 16.789 | 1 |
| 15 | Mast cells | Restf/f PR+/Cre | 32 | 36 | -10.817 | 1 |
| 16 | Neurons | Control | 10 | 22 | -54.428 | 0.039758708 |
| 16 | Neurons | Restf/f PR+/Cre | 46 | 34 | 35.068 | 0.039758708 |
| 17 | Fibroblasts | Control | 52 | 21 | 145.753 | 0 |
| 17 | Fibroblasts | Restf/f PR+/Cre | 2 | 33 | -93.910 | 0 |
| 18 | NKT | Control | 22 | 13 | 65.132 | 0.08602375 |
| 18 | NKT | Restf/f PR+/Cre | 12 | 21 | -41.966 | 0.08602375 |

**Supplemental Figure 7. Loss of REST leads to changes in uterine cell clusters. (A)** SingleR program predictions for cell clusters present in control and Rest$^{f/f}$ PR$^{+/Cre}$ mice uteri shown by a t-SNE plot. Cell clusters were given a number between 0-18. **(B)** Individual t-SNE plots for control and Rest$^{f/f}$ PR$^{+/Cre}$ mice showing changes in cell clusters. **(C)** Table showing expected number of cells in each cluster compared to actual number of cells in each cluster. Based on marker gene expression, cluster 0 (identified as fibroblasts) also represents cells of myometrial lineage. Percentage deviation and Bonferroni p-value show significant changes in clusters caused by loss of REST in the Rest$^{f/f}$ PR$^{+/Cre}$ mice.

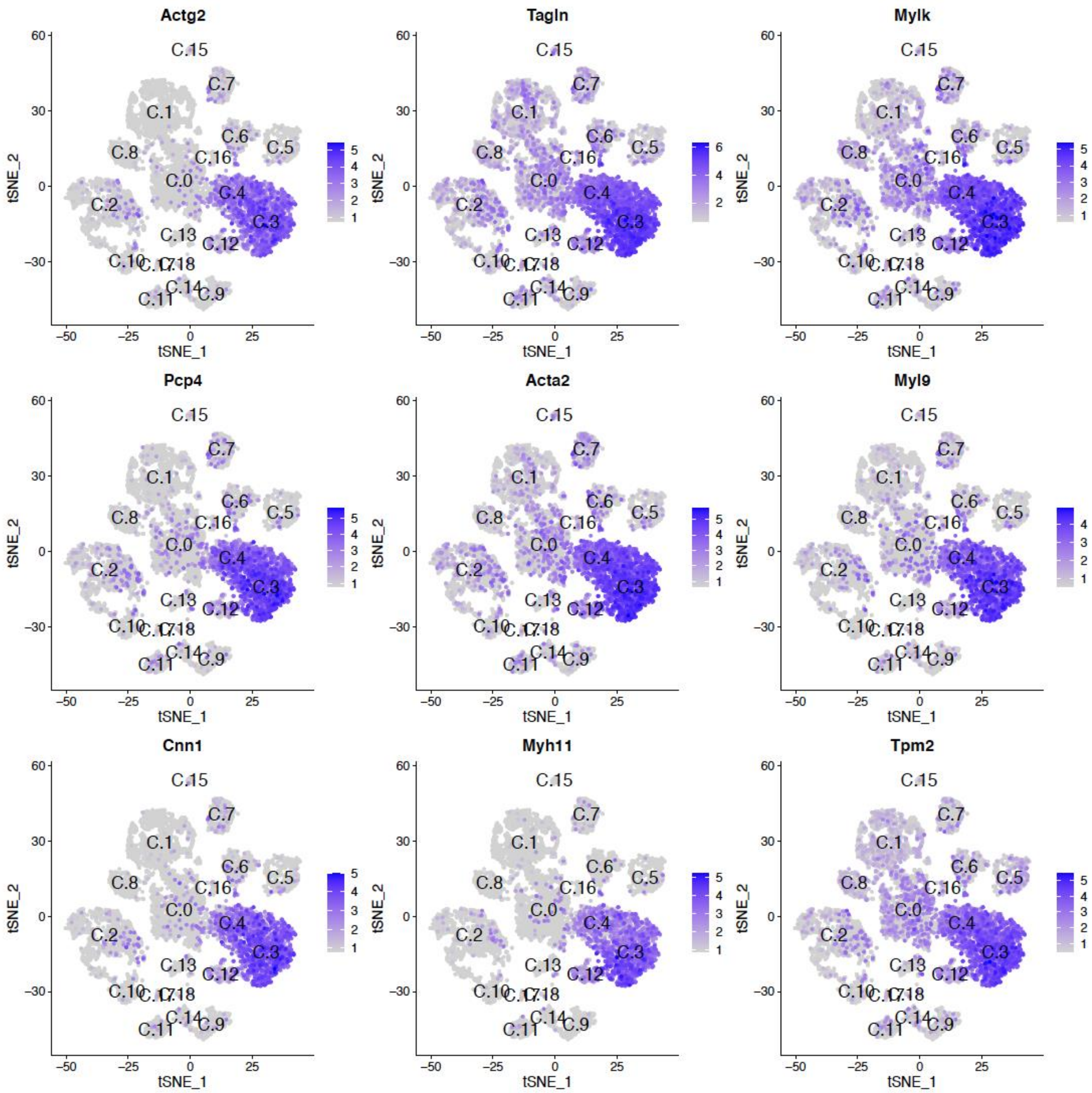


**Supplemental Figure 8. Clusters of cells showing conserved markers of smooth muscle cells**

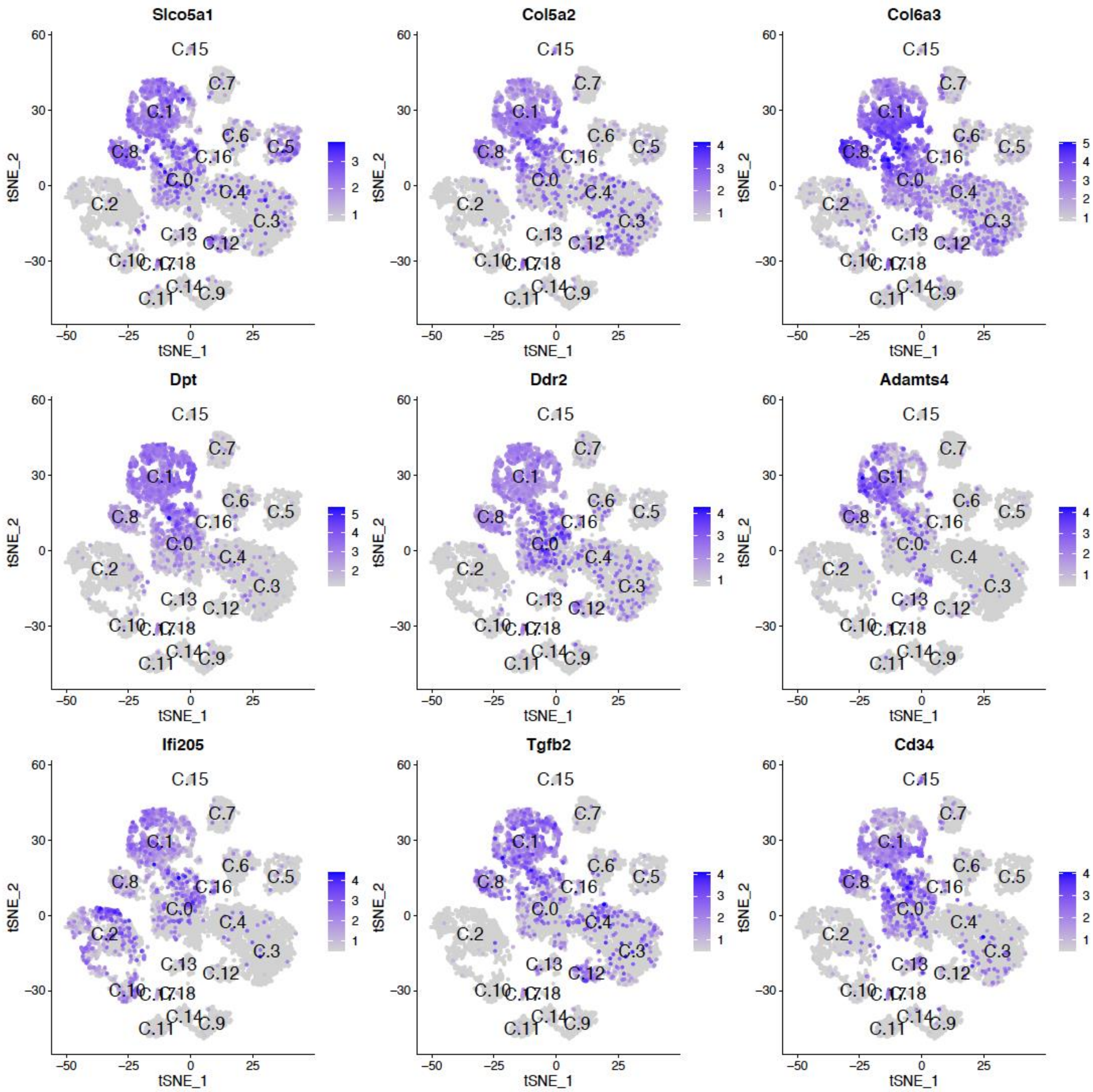


**Supplemental Figure 9. Clusters of cells showing conserved markers of uterine stromal cells, myometrial cells, and fibroblasts**

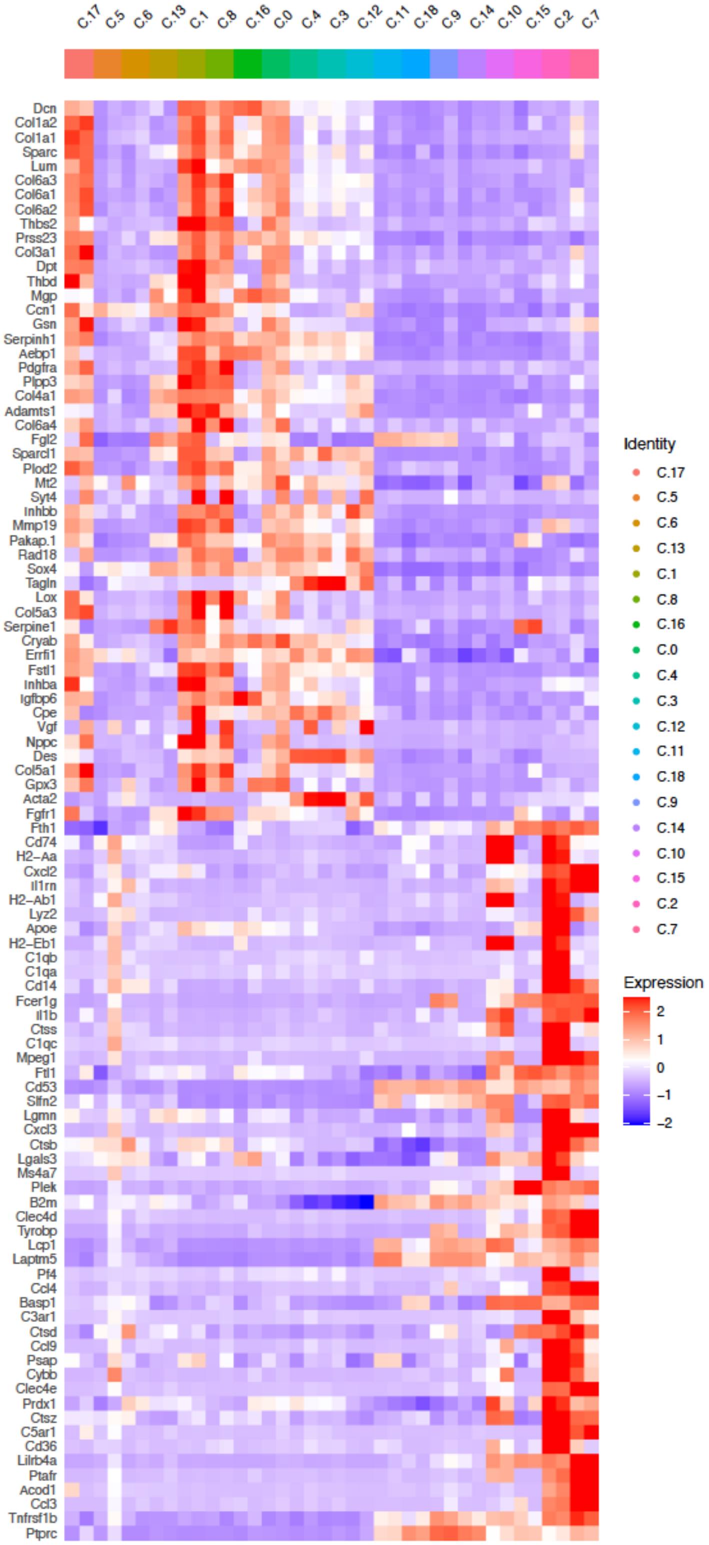


**Supplemental Figure 10. Top TCA features (average cluster expression) of genes**
Heatmap showing upregulation of ECM components in myometrial and stromal fibroblasts as well as smooth muscle cells (clusters 0, 1, 2, 3, 4, 8, and 17) in Rest$^{f/f}$ PR$^{+/Cre}$ mice.

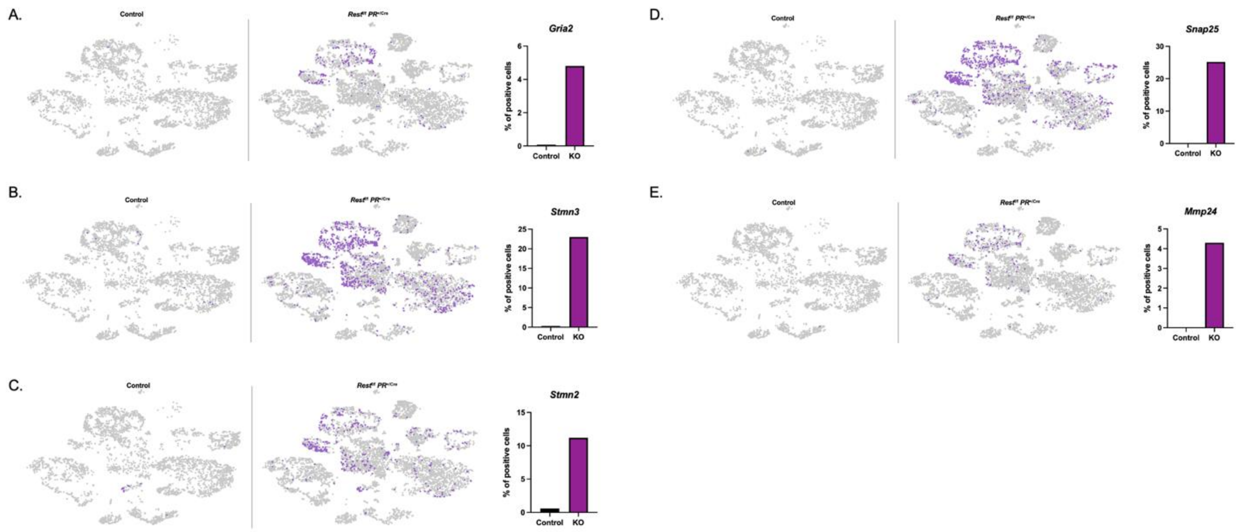


**Supplemental Figure 11. Expression of REST target genes in** *Rest*$^{f/f}$ *PR*$^{+/Cre}$ cKO mice**.** Comparative t-SNE plots of control and *Rest*$^{f/f}$ *PR*$^{+/Cre}$ cKO mice from single cell RNA sequencing showing expression (log 2-fold expression > 0) of REST target genes **(A)** *Gria2*, **(B)** *Stmn3*, and **(C)** *Stmn2*. Comparative t-SNE plots of control and *Rest*$^{f/f}$ *PR*$^{+/Cre}$ cKO mice uteri from single cell RNA sequencing showing expression (log 2-fold expression > 0) of Erα**-**REST target genes **(D)** *Snap25* and **(E)** *Mmp24*. Quantitative analysis (right panels) of the % of positive cells expressing REST targets, normalized using corresponding input cell numbers.

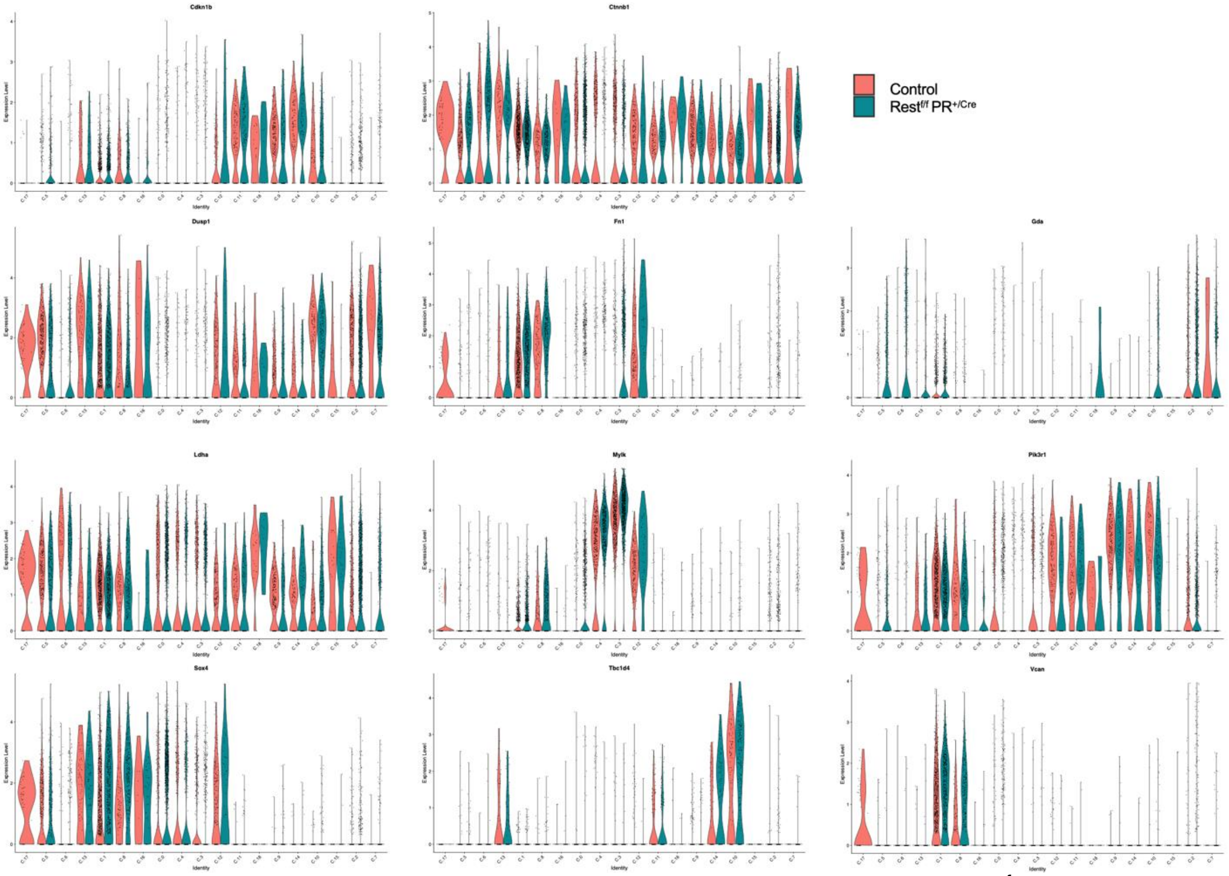


**Supplemental Figure 12. Cluster expression of REST target genes in** *Rest*$^{f/f}$ *PR*$^{+/Cre}$ cKO mice**.** Violin plots from single cell RNA sequencing showing expression (log 2-fold expression > 0) of REST target genes in different cell type clusters. X-axis represents the cluster number and Y-axis represents the gene expression level.

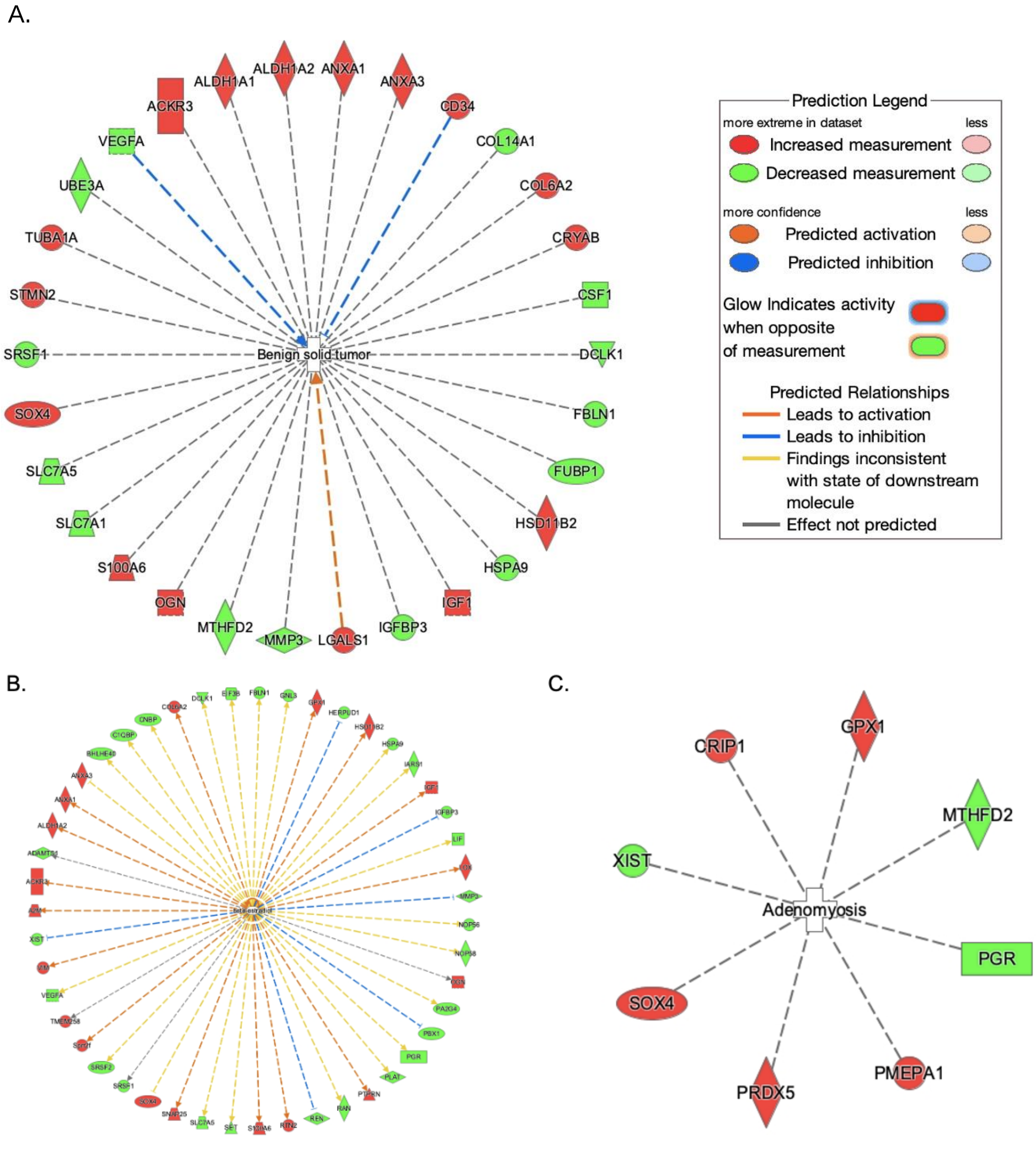


**Supplemental Figure 13. Ingenuity pathway analysis of cluster 0 in *Rest$^{f/f}$ PR$^{+/Cre}$* cKO mice. (A)** Gene network analysis of benign, solid tumor-associated genes per the IPA. **(B)** Gene network analysis of estrogen signaling-associated genes. **(C)** Gene network analysis of adenomyosis associated genes. GSE178141 cluster 0 genes which were significantly ($p < 0.05$) upregulated (red) or down regulated (green) in 5-month-old *Rest$^{f/f}$ PR$^{+/Cre}$* cKO mice compared to control.

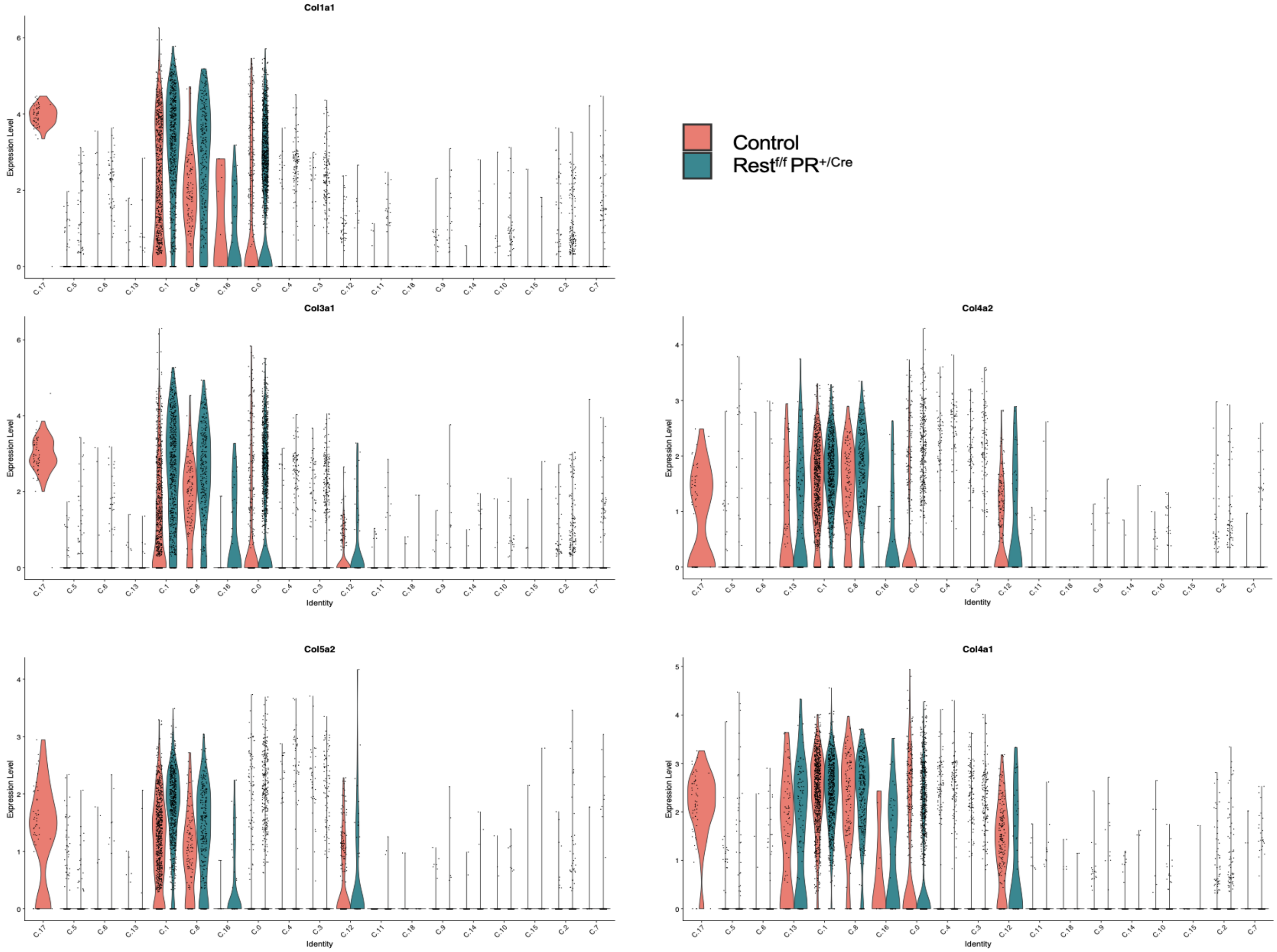


**Supplemental Figure 14. Cluster expression of collagen genes.** Violin plots showing control and *Rest*$^{f/f}$ *PR*$^{+/Cre}$ cKO mice from single cell RNA sequencing showing expression (log 2-fold expression > 0) of collagen genes in different cell type clusters. X-axis represents the cluster number and Y-axis represents the gene expression level.

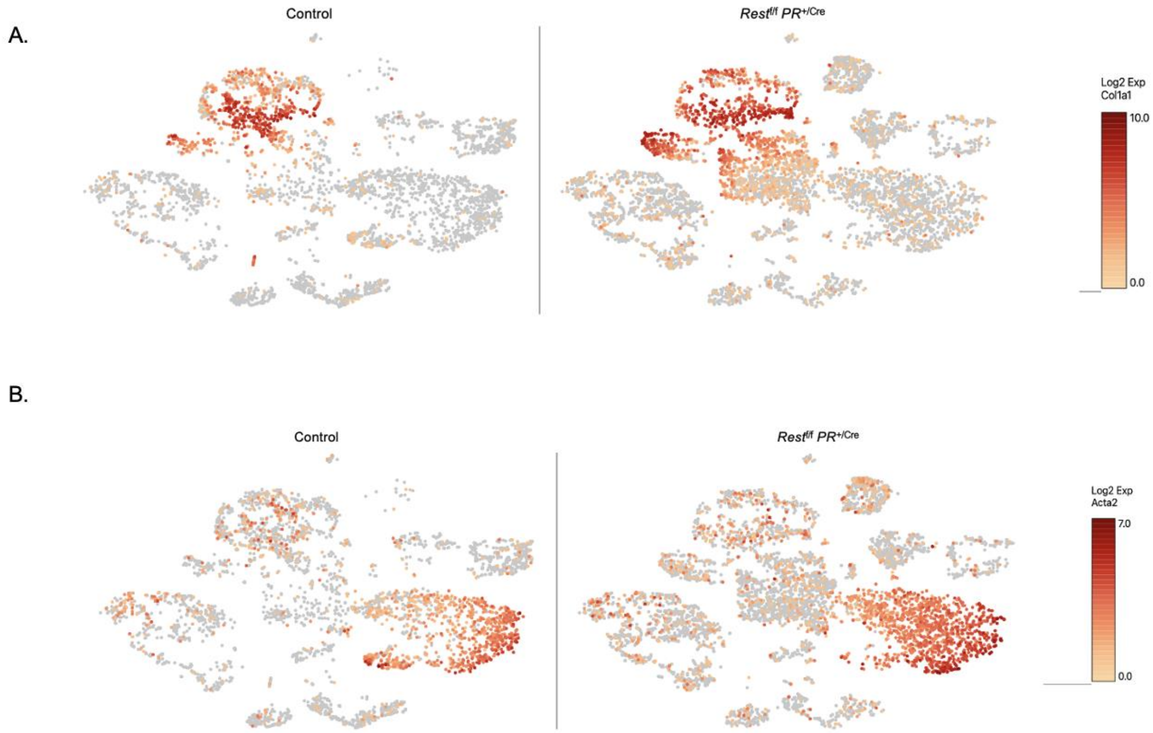


**Supplemental Figure 15. Single cell RNA sequencing analysis of ECM components in** *Rest*$^{f/f}$ *PR*$^{+/Cre}$ **cKO mice.** Comparative t-SNE plots showing expression (log 2-fold expression > 0) of extracellular matrix (ECM) components **(A)** *Col1a1*, and **(B)** *Acta2*. Dark red indicates higher levels of expression.

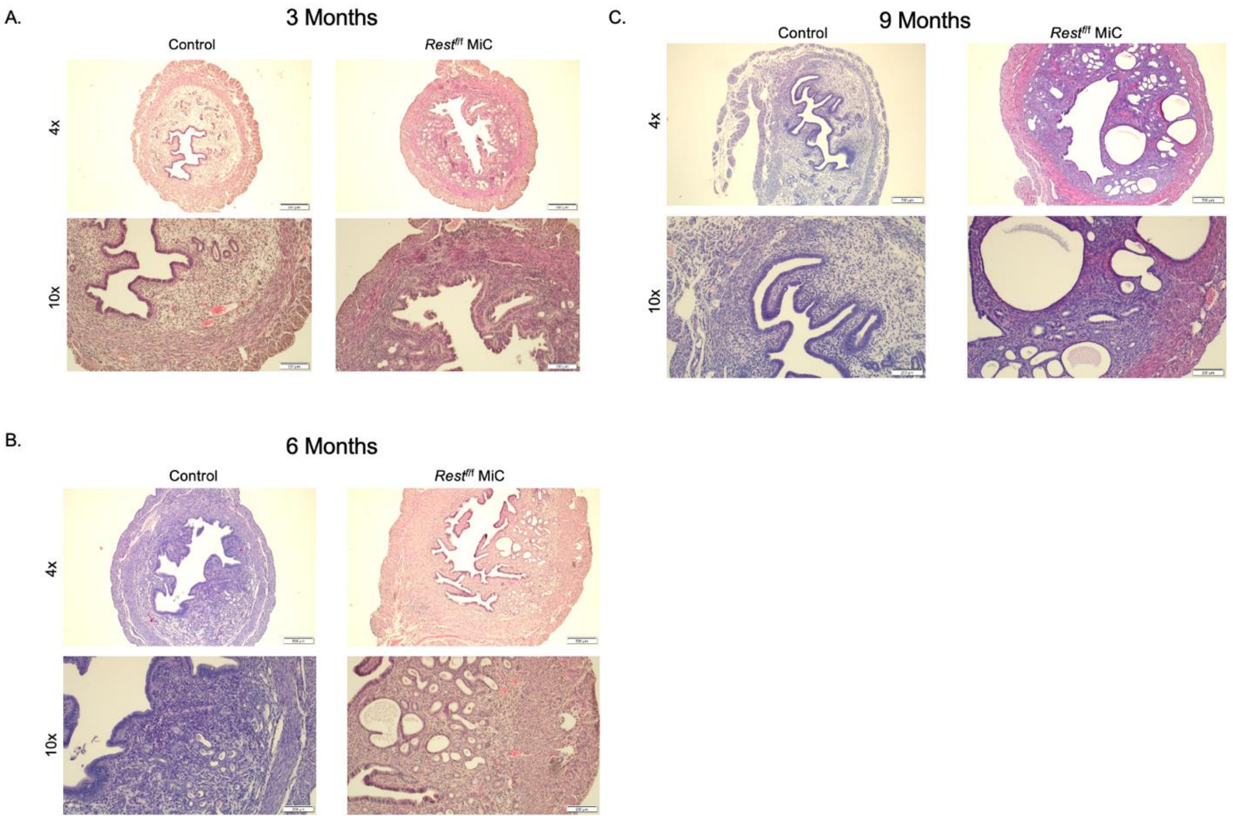


**Supplemental Figure 16. Characterization of *Rest*$^{f/f}$ MiC mouse uteri.**
H&E staining of uteri from control and *Rest*$^{f/f}$ MiC mice at (A) 3 months, **(B)** 6 months, and **(C)** 9 months in diestrus.

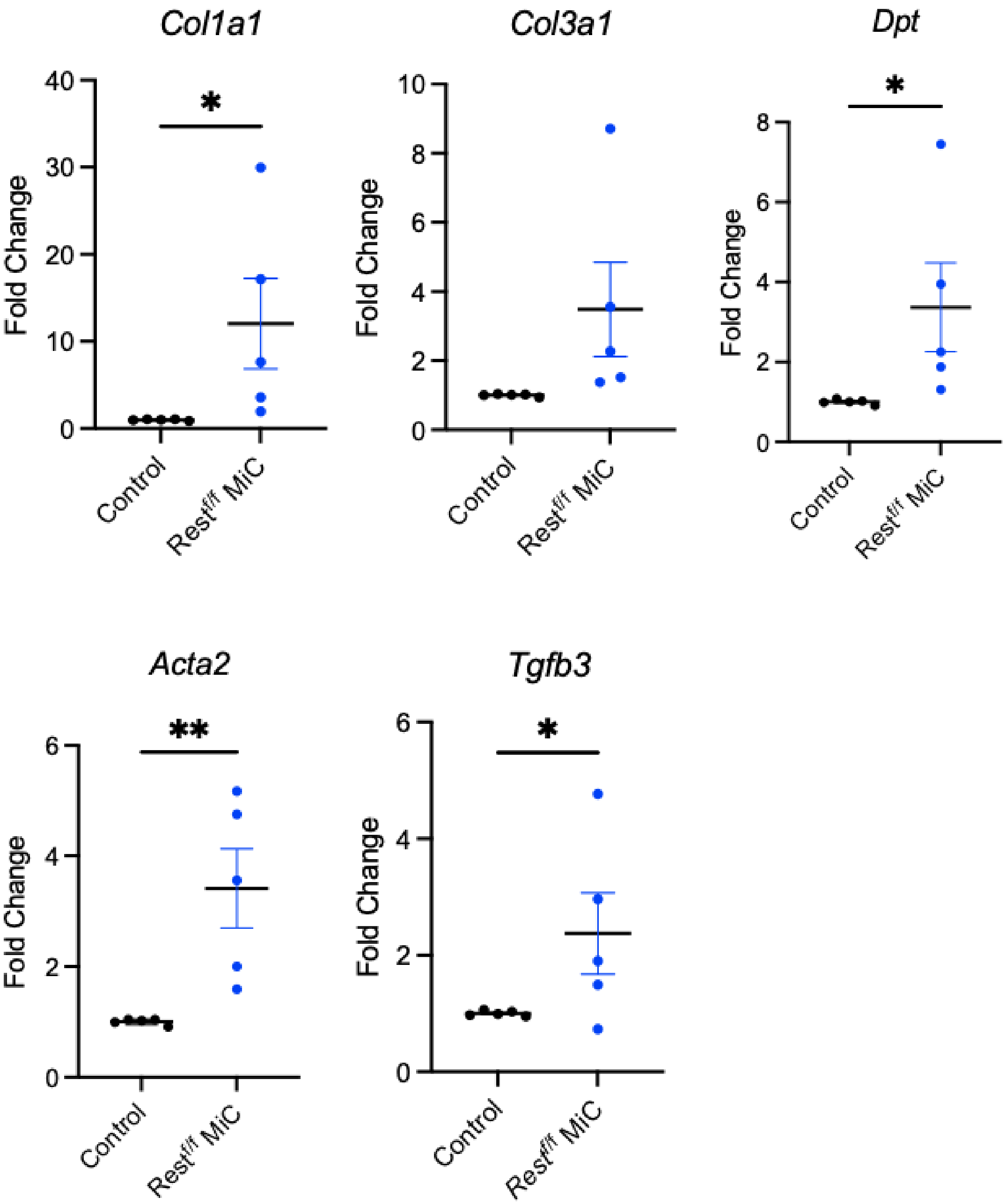


**Supplemental Figure 17. Increased expression of UL-specific genes in *Rest$^{f/f}$* MiC mice.** Quantitative RT-qPCR analysis of gene expression in uteri of 6-month-old *Rest$^{f/f}$* MiC and control mice (n=5). Error bars represent +SEM. Student's nonparametric T-test was performed, **P*<0.05, ***P*<0.01.

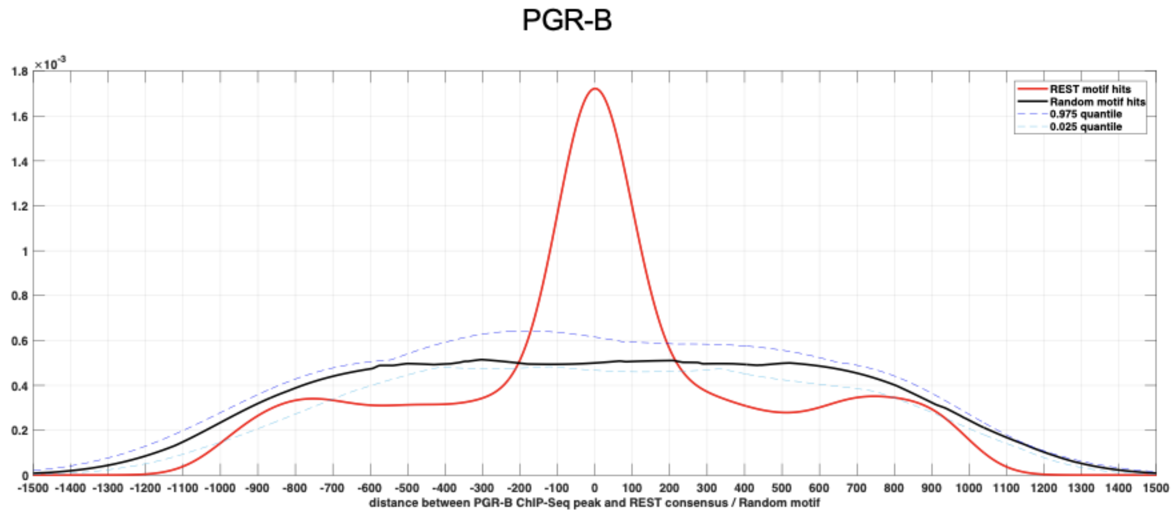


**Supplemental Figure 18. Interaction of REST with progesterone receptor B (PGR-B).** A conserved frequency of occurrence of REST binding sites (0) within 300 base pairs of PGR-B binding sites. X-axis represents distance in base pairs and y-axis represents density.

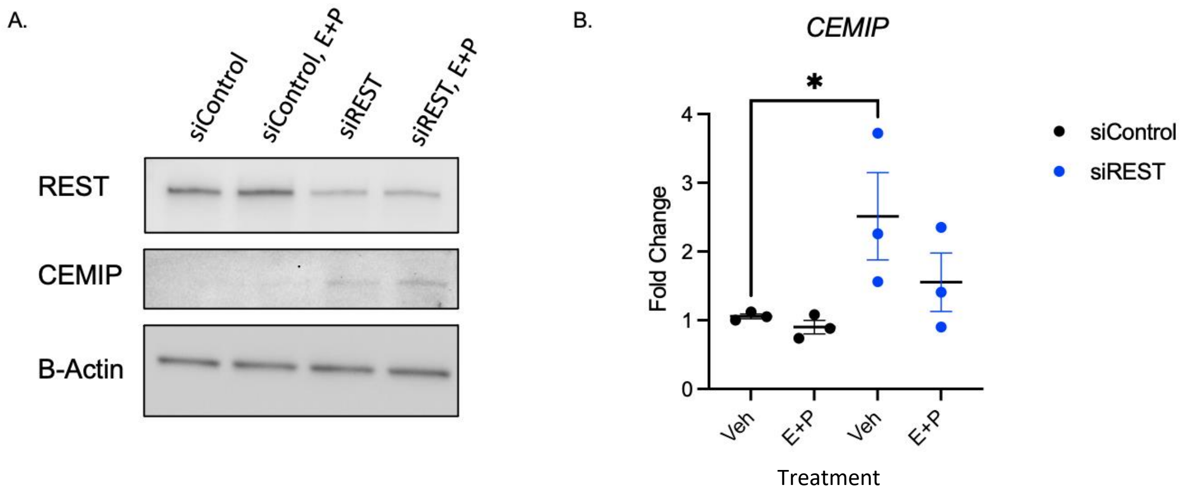


**Supplemental Figure 19. Loss of REST leads to upregulation of *CEMIP* in human primary myometrial cells. (A)** Western blot analysis showing REST and CEMIP expression in primary myometrial cells treated with siRNA targeting REST (siREST) or siRNA non-targeting (siControl). Cells were treated for 12 hours with either vehicle (PBS) or estrogen (E) and progesterone (P). β-actin was used as a loading control. **(B)** TaqMan RT-qPCR of *CEMIP* in primary myometrial cells treated for 24 hours with siRNA targeting REST (siREST) or siRNA Non-targeting (siControl). Cells were treated with either vehicle (PBS) or estrogen (E) and progesterone (P) (n=3). Error bars represent ±SEM. Student's T-test was performed,**P*=0.04

**Supplemental Table 1. Dysregulated REST target genes in human uterine leiomyoma specimens[a] .**

| Gene | Fold Change | Dysregulated in *Rest*$^{fl/fl}$ *Amhr2*$^{+/cre}$ cKO mice | Published role in uterus |
|---|---|---|---|
| DCX | 12.9723 | Yes | (15), (16) |
| ADAM12 | 5.11694 | No | (17) |
| KIF5C | 4.17796 | Yes | (15), (18) |
| SATB2 | 4.14309 | No | (19) |
| MEST | 4.08508 | No | (15), (16), (20) |
| MMP11 | 4.00553 | No | (21), (22) |
| IL17B | 3.86768 | Yes | (23) |
| PRLHR | 3.70861 | Yes | (5) |
| TYMS | 3.25197 | Yes | (16) |
| ANO1 | 2.95924 | No | (24) |
| CXCL13 | 2.85754 | No | (25) |
| RAD51B | 2.78883 | N/A | (26), (27) |
| PTHLH | 2.68783 | No | (28) |
| CKS2 | 2.60145 | Yes | (29) |
| DKK2 | 2.59632 | Yes | (30), (31) |
| GDF15 | 2.42375 | Yes | (32), (33) |
| PAK3 | 2.30551 | No | (34) |
| NAV2 | 2.29687 | No | (15) |
| TGFB2 | 2.27803 | No | (35) |
| EPHB1 | 2.23951 | Yes | (36) |
| SLC24A3 | 2.06676 | No | (37) |
| COL1A1 | 2.02989 | Yes | (38), (39) |
| SOX4 | 2.02886 | No | (40) |
| CDH2 | 2.0113 | No | (41) |
| CADM1 | 2.00376 | No | (42) |
| STRA6 | 1.9977 | Yes | (43) |
| RGS4 | 1.99677 | Yes | (44) |
| CYP1B1 | 1.96478 | Yes | (45) |
| VASH2 | 1.94743 | No | (46) |
| CCND1 | 1.92719 | Yes | (47) |
| NTM | 1.91407 | Yes | (48) |
| NRP2 | 1.90128 | No | (49) |
| PAPPA2 | 1.89855 | Yes | (50) |
| ITGA9 | 1.87332 | No | (51) |
| TRPS1 | 1.8703 | No | (52) |
| COL4A2 | 1.86263 | No | (15) |
| SEMA5A | 1.85369 | Yes | (53) |
| SDC1 | 1.84821 | No | (54), (55) |
| COL5A2 | 1.82204 | No | (56) |
| STMN2 | 1.81666 | Yes | (5) |
| VCAN | 1.79811 | No | (57), (58) |
| HMGA2 | 1.7872 | n/a | (59) |
| PDGFC | 1.75914 | No | (60) |
| SLC5A3 | 1.75614 | No | (61) |
| COL4A1 | 1.71354 | No | (62) |

| | | | |
|---|---|---|---|
| TET1 | 1.71152 | No | (63) |
| RRM2 | 1.70362 | No | (64) |
| GRIN2A | 1.70265 | Yes | (5) |
| TOP2A | 1.69869 | No | (65) |
| CTHRC1 | 1.67571 | No | (66) |
| CRMP1 | 1.65765 | Yes | (16) |
| PCNA | 1.64384 | Yes | (67) |
| PRLR | 1.64049 | Yes | (68) |
| MYLK | 1.63912 | No | (69) |
| MMP9 | 1.61055 | Yes | (70) |
| FAS | 1.59494 | No | (71) |
| CTNNB1 | 1.57665 | No | (72) |
| PCBP3 | 1.57286 | Yes | (73) |
| COL3A1 | 1.57017 | No | (74), (75) |
| PRKCB | 1.5695 | No | (36) |
| ASTN2 | 1.56375 | No | (76) |
| LHFPL3 | 1.56223 | N/A | (77) |
| MDM2 | 1.54577 | No | (78), (79) |
| IRS1 | 1.53128 | No | (80) |
| FN1 | 1.52285 | No | (40) |

[a] Fold change per **the** GEO dataset GSE13319

**Supplemental Table 2. REST ChIP-sequencing datasets**

| ENCODE Experiments for REST |
|---|
| ENCFF986RRJ |
| ENCFF896RCP |
| ENCFF796YFZ |
| ENCFF779CWH |
| ENCFF713ZPE |
| ENCFF706DRE |
| ENCFF669XCW |
| ENCFF668YET |
| ENCFF540FXB |
| ENCFF403CAJ |
| ENCFF313CII |
| ENCFF290ESJ |
| ENCFF274BBE |
| ENCFF208NUB |
| ENCFF107EWI |
| ENCFF023ZUW |

**Supplemental Table 3. Serum levels of estrogen and progesterone in *Rest$^{f/f}$ Amhr2$^{+/Cre}$* cKO mice during diestrus.**

| Sample | Estrogen (pg/mL) | Progesterone (ng/mL) |
|---|---|---|
| Wild type | 0.9 | 1.41 |
| Wild type | 1.2 | 1.90 |
| *Rest$^{f/f}$ Amhr2$^{+/Cre}$* | 2.9 | 2.53 |
| *Rest$^{f/f}$ Amhr2$^{+/Cre}$* | 2.1 | 2.44 |
| *Rest$^{f/f}$ Amhr2$^{+/Cre}$* | 0.5 | 26.92 |

**Supplemental Table 4. Dysregulated REST-PR target genes in *Rest*$^{f/f}$ *Amhr2*$^{+/Cre}$ cKO mouse uteri**[a]

| Gene | Fold Change |
|---|---|
| Adra1b | 2.21 |
| Angpt4 | 2.37 |
| Angptl1 | 2.80 |
| Apod | -2.62 |
| Bdnf | -2.04 |
| Blk | 1.75 |
| Bmp8a | 1.98 |
| Cacng2 | 65.70 |
| Ccbe1 | -2.44 |
| Ccnd1 | 1.56 |
| Cd55 | -2.13 |
| Cdh13 | -1.59 |
| Cdh4 | -2.06 |
| Cdk6 | -1.57 |
| Cdkn1b | -1.96 |
| Cdkn2b | -1.50 |
| Chat | 8.35 |
| Cisd1 | 1.60 |
| Cnr1 | 1.66 |
| Col1a1 | -1.92 |
| Col4a3 | -1.91 |
| Cyp19a1 | 3.23 |
| Drd3 | 30.30 |
| E2F2 | 1.52 |
| Edn2 | 2.49 |
| Epas1 | -1.64 |
| Ephb1 | -2.36 |
| Eps8 | -1.52 |
| Erh | 1.57 |
| Fibin | -2.74 |
| Gda | -1.57 |
| Gdf7 | 1.76 |
| Gprin1 | 13.98 |
| Grid1 | 2.35 |
| Grin2a | 1.52 |
| Habp2 | -6.38 |
| Hcst | 2.37 |
| Hpse2 | -1.70 |
| Htr1a | 3.67 |
| Il33 | -1.84 |
| Inmt | -5.57 |
| Itpr1 | -1.62 |
| Ldha | 1.61 |
| Lin28a | 9.19 |
| Mlph | 1.63 |
| Mmp24 | 59.61 |

| | |
|---|---|
| Mtus1 | -1.78 |
| Muc13 | -1.98 |
| Nek7 | -1.84 |
| Nppa | 24.35 |
| Nqo1 | 2.0 |
| Pdyn | -5.63 |
| Ppp2r2c | 1.69 |
| Psca | 1.87 |
| Reln | -1.61 |
| Rxfp3 | 2.52 |
| S100a8 | 15.44 |
| Scn2a1 | -1.72 |
| Smad5 | -1.80 |
| Snap25 | 24.15 |
| Stmn2 | 2.96 |
| Stra6 | 1.58 |
| Synpr | 2.45 |
| Wnt9b | 1.51 |
| Xkr7 | 111.56 |

[a] Fold change per the GEO data set GSE178141

**Supplemental Table 5. Dysregulated REST-PR target genes in human uterine leiomyoma specimens[a]**

| Gene | p-value | Fold Change |
|---|---|---|
| ADAM12 | 0.0001 | 5.11 |
| CDKN2B | 0.0143 | -1.50 |
| CEMIP | 3.23 E-06 | 5.79 |
| CNR1 | 0.0003 | -1.53 |
| COL1A1 | 1.36E-05 | 2.03 |
| DUSP1 | 2.90E-05 | -2.67 |
| EPHB1 | 0.0014 | 2.24 |
| GPM6B | 0.0008 | 2.19 |
| GRP | 0.0691 | 1.83 |
| IL6R | 0.0002 | -1.51 |
| IRS1 | 0.0046 | 1.53 |
| KCNMA1 | 0.0055 | 1.71 |
| LHFPL3 | 0.0387 | 1.56 |
| MTUS1 | 0.0002 | -1.78 |
| MYLK | 0.0079 | 1.64 |
| NTM | 0.0022 | 1.91 |
| PIK3R1 | 0.0009 | 1.78 |
| PTCH1 | 0.0001 | -1.95 |
| SATB2 | 1.77E-07 | 4.14 |
| SEMA5A | 9.18E-06 | 1.85 |
| SRD5A1 | 0.0001 | 1.62 |
| STRA6 | 0.0006 | 2.00 |
| TNN | 0.0027 | 1.82 |
| TRPC6 | 1.01E-05 | 2.07 |

[a] Fold change per the GEO data set GSE13319
